## Supplemental materials for "*Streptomyces* alleviate abiotic stress in plant by producing pteridic acids"

### Table of Contents


| Content | Page |
|---|---|
| **Tab. 1**. The biosynthetic gene clusters in *S. iranensis*. | 3 |
| **Tab. 2**. The annotation of pteridic acids core biosynthetic genes in *S. iranensis*. | 5 |
| **Tab. 3**. Alignment of conserved motifs in the active site of AT domains. | 6 |
| **Tab. 4**. The list of *Streptomyces* strains that can produce pteridic acids, elaiophylin or harboring *pta* gene cluster. | 7 |
| **Tab. 5**. Summary of strains and plasmids used in this study. | 9 |
| **Tab. 6**. $^1$H (800 MHz) NMR data for **1**–**2** (in MeOD). | 10 |
| **Tab. 7**. $^{13}$C (200 MHz) NMR data for **1**–**2** (in MeOD). | 11 |
| **Tab. 8**. Summary of primers used in this study. | 12 |
| **Fig. 1**. Enrichment evaluation of *S. iranensis* in rhizosphere soil. | 13 |
| **Fig. 2**. Barley experiments of *S. iranensis* and *S. iranensis*/Δ*ptaA* treatment. | 13 |
| **Fig. 3**. Mirror plots comparing spectra from known metabolites from *S. iranensis* to standard spectra deposited in GNPS. | 14 |
| **Fig. 4**. $^1$H NMR spectrum of **1.** | 15 |
| **Fig. 5**. $^{13}$C NMR spectrum of **1.** | 15 |
| **Fig. 6**. NOESY spectrum of **1**. | 16 |
| **Fig. 7**. COSY spectrum of **1**. | 16 |
| **Fig. 8**. HSQC spectrum of **1**. | 17 |
| **Fig. 9**. HMBC spectrum of **1**. | 17 |
| **Fig. 10**. Mass spectrum of **1**. | 18 |
| **Fig. 11**. Selected HMBC correlations for **1** and **2**. | 18 |
| **Fig. 12**. Crystal structure of **1**. | 18 |
| **Fig. 13**. $^1$H NMR spectrum of **2**. | 19 |
| **Fig. 14**. $^{13}$C NMR spectrum of **2**. | 19 |
| **Fig. 15**. COSY spectrum of **2**. | 20 |
| **Fig. 16**. NOESY spectrum of **2**. | 20 |
| **Fig. 17**. H2BC spectrum of compound **2**. | 21 |
| **Fig. 18**. HSQC spectrum of **2**. | 21 |
| **Fig. 19**. HMBC spectrum of **2**. | 22 |
| **Fig. 20**. Mass spectrum of **2**. | 22 |
| **Fig. 21**. Primary root length of *Arabidopsis* seedling treated with different concentrations of pteridic acids H and F. | 23 |
| **Fig. 22**. Kidney beans growth experiment with pure pteridic acids. | 23 |
| **Fig. 23**. Pteridic acid H and ABA at 1 ng $mL^{-1}$ help Mung beans against heavy metal stress. | 24 |
| **Fig. 24**. Multiple sequence alignment of KR domains. | 25 |
| **Fig. 25**. Multiple sequence alignment of DH domains. | 26 |
| **Fig. 26**. Stability test of pteridic acids. | 27 |
| **Fig. 27**. The schematic of optimized genetic manipulation in *S. iranensis* by using CRISPR-cBEST system. | 28 |
| **Fig. 28**. Complementation experiment of ptaA-inactivation mutant of *S. iranensis*. | 29 |
| **Fig. 29**. Multiple sequence alignment of TE domains. | 30 |
| **Fig. 30**. The HR-LC-MS/MS analysis of pteridic acids in *S. violaceusniger* Tu 4113 and *S. rapamycinicus* NRRL 5491. | 31 |
| **Fig. 31**. The phylogenetic analysis of potential pteridic acids *Streptomyces* producers. | 32 |
| **Fig. 32**. Genome similarity analysis based on the alignment of 15 sequenced *pta*-containing *Streptomyces* genomes. | 33 |
| **Fig. 33**. The genome synteny analysis of 15 *pta*-containing *Streptomyces* strains. | 34 |
| **Fig. 34**. Identification of pteridic acids and elaiophylin in *S. albus* DSM 41398. | 35 |
| **Fig. 35**. The secondary metabolites biosynthetic gene clusters profiles of 15 *pta*-containing *Streptomyces* strains with complete genome information. | 36 |
| **Fig. 36**. The secondary metabolites BGCs similarity network of 15 pta-containing *Streptomyces* strains with complete genome information. | 36 |
| **References** | 37 |

**Tab. 1.** The biosynthetic gene clusters in *S. iranensis*.

| Cluster | Type | Position | | Most similar known cluster | Similarity |
|---|---|---|---|---|---|
| Cluster 1 | lassopeptide | 47,826 | 67,848 | SSV-2083 | 18% |
| Cluster 2 | NRPS-like, T1PKS, NRPS | 89,243 | 264,517 | argimycins | 10% |
| Cluster 3 | terpene, NRPS | 310,755 | 369,526 | carotenoid | 63% |
| Cluster 4 | NRPS | 584,615 | 633,798 | coelichelin | 100% |
| Cluster 5 | butyrolactone | 920,071 | 929,026 | cyphomycin | 9% |
| Cluster 6 | phosphonate, acyl_amino_acids, butyrolactone, NRPS-like, T1PKS, hserlactone | 983,576 | 1,129,298 | lydicamycin | 20% |
| Cluster 7 | T1PKS | 1,216,733 | 1,353,478 | azalomycin F3a | 100% |
| Cluster 8 | T1PKS | 1,525,866 | 1,708,371 | nigericin | 100% |
| Cluster 9 | T1PKS | 1,812,276 | 1,890,013 | elaiophylin | 87% |
| Cluster 10 | redox-cofactor | 1,914,682 | 1,936,746 | | |
| Cluster 11 | hserlactone | 2,054,025 | 2,074,780 | daptomycin | 4% |
| Cluster 12 | butyrolactone | 2,085,494 | 2,096,426 | | |
| Cluster 13 | T1PKS, NRPS | 2,274,879 | 2,327,099 | meilingmycin | |
| Cluster 14 | NRPS, T3PKS, other | 2,383,216 | 2,482,446 | feglymycin | |
| Cluster 15 | NRPS, T1PKS | 2,535,309 | 2,739,409 | α-lipomycin | |
| Cluster 16 | terpene | 2,985,464 | 3,007,479 | hopene | 76% |
| Cluster 17 | T1PKS | 3,153,626 | 3,235,531 | macbecin / macbecin II | 34% |
| Cluster 18 | T2PKS | 3,304,807 | 3,374,688 | spore pigment | 83% |
| Cluster 19 | T1PKS | 3,499,162 | 3,559,357 | s56-p1 | 11% |
| Cluster 20 | RiPP-like | 3,612,469 | 3,622,637 | | |
| Cluster 21 | siderophore | 3,792,665 | 3,803,134 | | |
| Cluster 22 | T2PKS | 4,140,701 | 4,213,204 | isoindolinomycin | 61% |
| Cluster 23 | NRPS-like | 4,442,739 | 4,484,596 | echosides | 100% |
| Cluster 24 | siderophore | 5,229,788 | 5,240,857 | desferrioxamin B | 100% |
| Cluster 25 | terpene | 6,441,503 | 6,461,832 | geosmin | 100% |
| Cluster 26 | ladderane, arylpolyene, NRPS | 6,705,172 | 6,807,080 | atratumycin | 57% |
| Cluster 27 | NRPS | 7,345,535 | 7,387,933 | ochronotic pigment | 75% |
| Cluster 28 | ladderane | 7,449,836 | 7,489,866 | atratumycin | 34% |
| Cluster 29 | RRE-containing | 7,510,172 | 7,530,691 | granaticin | 10% |
| Cluster 30 | T1PKS | 7,664,407 | 7,848,116 | mediomycin A | 68% |
| Cluster 31 | terpene | 8,396,931 | 8,416,950 | | |
| Cluster 32 | ectoine | 9,062,415 | 9,072,819 | ectoine | 100% |
| Cluster 33 | siderophore | 9,237,702 | 9,251,453 | | |
| Cluster 34 | terpene | 9,301,372 | 9,319,391 | BE-43547s | 25% |
| Cluster 35 | nucleoside | 9,535,814 | 9,572,869 | toyocamycin | 40% |
| Cluster 36 | nucleoside, NRPS, T1PKS, NRPS-like | 9,677,504 | 9,821,457 | rapamycin | 82% |
| Cluster 37 | PKS-like | 9,909,290 | 9,950,318 | rustmicin | 33% |
| Cluster 38 | terpene | 10,000,500 | 10,020,321 | 2-methylisoborneol | 100% |
| Cluster 39 | terpene | 10,464,893 | 10,484,016 | pristinol | 100% |
| Cluster 40 | T1PKS, ladderane,arylpolyene | 10,578,295 | 10,629,355 | atratumycin | 28% |

| | | | | | |
|---|---|---|---|---|---|
| Cluster 41 | T1PKS, NRPS-like | 10,835,982 | 10,918,034 | hygrocins | 83% |
| Cluster 42 | lanthipeptide-class-ii | 11,011,708 | 11,035,079 | reveromycin A | 9% |
| Cluster 43 | hglE-KS, T1PKS, RiPP-like | 11,062,970 | 11,124,110 | | |
| Cluster 44 | T1PKS, NRPS | 11,127,758 | 11,269,230 | X-14547 | 21% |
| Cluster 45 | terpene | 11,320,894 | 11,341,820 | brasilicardin A | 38% |
| Cluster 46 | terpene | 11,470,415 | 11,491,323 | | |
| Cluster 47 | T1PKS, NRPS-like | 11,577,719 | 11,624,306 | niphimycins C-E | 9% |
| Cluster 48 | betalactone | 11,818,784 | 11,847,595 | Sch-47554 / Sch-47555 | 7% |
| Cluster 49 | NRPS, NRPS-like, T1PKS | 12,040,448 | 12,123,871 | echosides | 11% |
| Cluster 50 | lassopeptide | 12,143,153 | 12,165,683 | SSV-2083 | 18% |

**Tab. 2**. The annotation of pteridic acids core biosynthetic genes in *S. iranensis*.

| ORF | Size[a] | Proposed functions | SI/ID[b] | Protein homologue and origin |
|---|---|---|---|---|
| *Pta5* | 1041 | 3-oxoacyl-ACP synthase III | 98/97 | WP_210951575.1, *Streptomyces* sp. MK37H |
| *pta4* | 572 | 3-hydroxyacyl-CoA dehydrogenase | 98/97 | WP_020866330.1, *Streptomyces rapamycinicus* |
| *pta3* | 958 | LuxR family transcriptional regulator | 99/99 | WP_020866329.1, *Streptomyces rapamycinicus* |
| *pta2* | 316 | glucose-1-phosphate thymidylyltransferase | 99/99 | RLV74943.1, *Streptomyces rapamycinicus* NRRL 5491 |
| *pta1* | 324 | TDP glucose 4,6 dehydratase | 98/98 | WP_214665150.1, *Streptomyces javensis* |
| *ptaA* | 4535 | Type I PKS | 90/88 | WP_214609358.1, *Streptomyces malaysiensis* |
| *ptaB* | 1746 | Type I PKS | 91/89 | WP_037957959.1, *Streptomyces* sp. PRh5 |
| *ptaC* | 1655 | Type I PKS | 95/94 | WP_138910801.1, *Streptomyces* sp. DASNCL29 |
| *ptaD* | 3395 | Type I PKS | 96/95 | WP_020866322.1, Streptomyces rapamycinicus |
| *ptaE* | 2112 | Type I PKS | 95/95 | WP_201848053.1, *Streptomyces* sp. 110 |
| *pta1** | 261 | Thioesterase | 97/95 | GDY58888.1, *Streptomyces violaceusniger* |
| *pta2** | 417 | Glycosyltransferase | 99/98 | MBP8534388.1, *Streptomyces* sp. MK37H |
| *pta3** | 196 | dTDP-4-dehydrorhamnose 3,5-epimerase | 97/97 | WP_210946413.1, *Streptomyces* sp. MK37H |
| *pta4** | 304 | Exporter (membrane domain) | 100/99 | WP_138910796.1, *Streptomyces* sp. DASNCL29 |
| *pta5** | 245 | Exporter (ATPase domain) | 99/99 | WP_020866316.1, *Streptomyces rapamycinicus* |
| *pta6** | 420 | Two-component regulator (sensor/kinase domain) | 97/97 | WP_199334865.1, *Streptomyces* sp. GMR22 |
| *pta7** | 223 | Two-component regulator (effector domain) | 98/98 | WP_138910795.1, *Streptomyces* sp. DASNCL29 |
| *pta8** | 321 | NAD(P)-dependent oxidoreductase | 96/96 | WP_164428021.1, *Streptomyces rhizosphaericus* |
| *pta9** | 328 | aldo/keto reductase | 98/98 | WP_191066610.1, *Streptomyces* sp. 5-10 |
| *pta10** | 469 | NDP-hexose 2,3 dehydratase | 97/95 | GDY58900.1, *Streptomyces violaceusniger* |
| *pta11** | 446 | Crotonyl-CoA reductase | 99/99 | WP_020866310.1, *Streptomyces rapamycinicus* |

a, The size of amino acids; b, similarity-identity ratio.

**Tab. 3**. Alignment of conserved motifs in the active site of AT domains.

| domain name | specificity | 198 | 199 | 200 | 201 |
|---|---|---|---|---|---|
| ave_AT5 | Malonyl-CoA | H | A | F | H |
| nid_AT3 | Malonyl-CoA | H | A | F | H |
| epo_AT2 | Malonyl-CoA | H | A | F | H |
| amp_AT18 | Malonyl-CoA | H | A | F | H |
| rif_AT2 | Malonyl-CoA | H | A | F | H |
| pta_LD | Malonyl-CoA | H | A | F | H |
| pta_AT2 | Malonyl-CoA | H | A | F | H |
| pta_AT6 | Malonyl-CoA | H | A | F | H |
| pta_AT7 | Malonyl-CoA | I | A | A | H |
| ave_AT1 | Methylmalony-CoA | Y | A | S | H |
| nid_AT4 | Methylmalony-CoA | Y | A | S | H |
| ery_AT4 | Methylmalony-CoA | Y | A | S | H |
| amp_AT2 | Methylmalony-CoA | Y | A | S | H |
| rif_AT7 | Methylmalony-CoA | Y | A | S | H |
| pta_AT3 | Methylmalony-CoA | Y | A | S | H |
| pta_AT4 | Methylmalony-CoA | Y | A | S | H |
| pta_AT5 | Methylmalony-CoA | Y | A | S | H |
| nid_AT5 | Ethylmalonyl-CoA | T | A | G | H |
| tyl_AT5 | Ethylmalonyl-CoA | T | A | G | H |
| pta_AT1 | Ethylmalonyl-CoA | T | A | G | H |
| ery_LD | Propionyl-CoA | M | A | A | H |
| meg_LD | Propionyl-CoA | M | A | A | H |
| epo_AT3 | Flexible | H | A | S | H |

* Abbreviations: LD, loading module; ave, avermectin; nid, niddamycin; epo, epothilone; amp, amphotericin; rif, rifamycin; ery, erythromycin; tyl, tylactone; meg, megalomicin; pta, pteridic acids.

**Tab. 4**. The *Streptomyces* strains that can produce elaiophylin, pteridic acids, or harbor *pta* BGC.

| NO | Strain name | Geographic location | Source | Reference/Assembly |
|---|---|---|---|---|
| 1 | *Streptomyces iranensis* HM 35 | Isfahan City, Iran | rhizosphere | This study |
| 2 | *Streptomyces* sp. 219807 | Sanya, Hainan, China | mangrove soil | 1 |
| 3 | *Streptomyces* sp. SCSGAA 0027 | South China Sea, China | gorgonian-associated | 2 |
| 4 | *Streptomyces* sp. 7-145 | Heishijiao Bay, Dalian, China | marine-sediment | 3 |
| 5 | *Streptomyces melanosporofaciens* | Italy | soil | 4 |
| 6 | *Streptomyces autolyticus* CGMCC 0516 | Yunnan, China | soil | 5 |
| 7 | *Streptomyces* sp. DSM 3816 | Kypcerissia, Greece | soil | 6 |
| 8 | *Streptomyces* sp. BCC 71188 | Nakhon Si Thammarat Province, Thailand | soil | 7 |
| 9 | *Streptomyces* sp. BCC 72023 | Chumphon province, Thailand | plant-associated | 8 |
| 10 | *Streptomyces* sp. BS 1261 | New Zealand | soil | 9 |
| 11 | *Streptomyces* sp. ICBB 9297 | Jatiroto, East Java, Indonesia | soil | 10 |
| 12 | *Streptomyces* sp. SNA-4606 | Towada-shi, Aomori Prefecture, Japan | soil | 11 |
| 13 | *Streptomyces* sp. MCY-846 | cheju-island, Korea | soil | 12 |
| 14 | *Streptomyees albiflaviniger* SCSIO ZJ28 | South China Sea, China | marine-sediment | 13 |
| 15 | *Streptomyces* sp. USC-16018 | Hastings Point, NSW, Australia | marine | 14 |
| 16 | *Streptomyces* sp. SPMA113 | Prajinburi Province, Thailand | soil | 15 |
| 17 | *Streptomyces* sp. LZ35 | Ji'mei, Xia'men, China | soil | 16 |
| 18 | *Streptomyces malaysiensis* DSM 4137 | Germany | soil | 17 |
| 19 | *Streptomyces* sp. BCa1 | Borra Caves, India | soil | 18 |
| 20 | *Streptomyces* sp. IFM11958 | Sakuragi cemetery, Chiba city, Japan | soil | 19 |
| 21 | *Streptomyces malaysiensis* OUCMDZ-2167 | South China Sea, China | marine | 20 |
| 22 | *Streptomyces* sp. 11-1-2 | Newfoundland, Canada | plant-associated | 21 |
| 23 | *Streptomyces* sp. HNM0561 | Hainan, China | marine-sediment | 22 |
| 24 | *Streptomyces* sp. GMR 22 | Wanagama Forest, Indonesia | soil | 23 |
| 25 | *Streptomyces* sp. NTK 935 | Canary Basin | marine sediment | 24 |
| 26 | *Streptomyces* sp. NTK 937 | Canary Basin | marine sediment | 24 |
| 27 | *Streptomyces* sp. RJA 2928 | Papua New Guinea | marine sediment | 25 |
| 28 | *Streptomyces rapamycinicus* NRRL 5491 | Easter island, Fiji | soil | 26 |
| 29 | *Streptomyces pseudouerticillus* YN 17707 | Xishuangbanna, Yunnan, China | soil | 27 |
| 30 | *Streptomyces* sp. SCSIO ZS0520 | Okinawa, Japan | marine sediment | 28 |
| 31 | *Streptomyces hygroscopicus* NND-52 | Suqian, Jiangsu, China | soil | 29 |
| 32 | *Streptomyces* hygroscopicus NO.662 | Sapporo-city, Hokkaido, Japan | soil | 30 |
| 33 | *Streptomyces hygroscopicus* TP-A0451 | Toyama, Japan | plant-associated | 31 |
| 34 | *Streptomyces hygroscopicus* CH-7 | Vojvodina, Serbia | soil | 32 |
| 35 | *Streptomyces hygroscopicus* 17997 | Yunnan, China | soil | 33 |
| 36 | *Streptomyces hygroscopicus* ACTMS-9H | Amazon, Brazil | rhizosphere | 34 |
| 37 | *Streptomyces hygroscopicus* XM 201 | Xiamen, Fujian, China | soil | 35 |
| 38 | *Streptomyces yatensis* DSM 41771 | New Caledonia | Ultramafic soil | 36 |
| 39 | *Streptomyces solisilvae* HNM0141 | Bawangling, Hainan, China | soil | 37 |
| 40 | *Streptomyces* sp. NA02950 | Hainan, China | marine-sediment | 38 |

| | | | | |
|---|---|---|---|---|
| 41 | *Streptomyces* sp. PRh5 | Dongxiang, China | plant-associated | 39 |
| 42 | *Streptomyces albus* DSM 41398 | Fuji City, Shizuoka Pref, Japan | soil | GCA_000827005.1 |
| 43 | *Streptomyces samsunensis* SA31 | Songkhla, Thailand | soil | GCA_013345665.1 |
| 44 | *Streptomyces* sp. MK37H | Antalya, Turkey | soil | GCA_018035285.1 |
| 45 | *Streptomyces* sp. 4503 | Guangxi, China | mangrove sediment | GCA_018883605.1 |
| 46 | *Streptomyces* sp. t39 | Austin, Texas, USA | soil | GCA_008042045.1 |
| 47 | *Streptomyces* sp. 5-10 | Hainan, China | plant-associated | GCA_014712245.1 |
| 48 | *Streptomyces antioxidans* MUSC164 | Malaysia | mangrove | GCA_000968685.2 |
| 49 | *Streptomyces malaysiensis* F913 | Chongqing, China | soil | GCA_002891865.1 |
| 50 | *Streptomyces* sp. DASNCL29 | Unkeshwar, India | soil | GCA_005938145.1 |
| 51 | *Streptomyces* sp. WAC05858 | Germeny | soil | GCA_003949695.1 |
| 52 | *Streptomyces rhizosphaericus* 0250 | Taian, China | soil | GCA_010892295.1 |
| 53 | *Streptomyces* sp. NEAU-YJ-81 | Harbin, China | soil | GCA_017592595.1 |
| 54 | *Streptomyces rhizosphaericus* NRRL B-24304 | Indonesia | rhizosphere | GCA_002155885.1 |
| 55 | *Streptomyces malaysiensis* TY049-057 | Bidor Perak, Malaysia | soil | GCA_008033485.1 |
| 56 | *Streptomyces malaysiensis* DSM 14702 | Germany | soil | GCA_011800555.1 |
| 57 | *Streptomyces cangkringensis* DSM 41769 | South Korea | - | GCA_019059395.1 |
| 58 | *Streptomyces endocoffeicus* CA3R110 | Lampang, Thailand | plant-associated | GCA_016741935.1 |
| 59 | *Streptomyces indonesiensis* DSM 41759 | Yogyakarta, Indonesia | rhizosphere | GCA_018138705.1 |
| 60 | *Streptomyces rhizosphaericus* DSM 41760 | Yogyakarta, Indonesia | rhizosphere | GCA_017942185.1 |
| 61 | *Streptomyces asiaticus* DSM 41761 | Yogyakarta, Indonesia | rhizosphere | GCA_018138715.1 |
| 62 | *Streptomyces* sp. RCU064 | Nong Jum Rung, Thailand | Peat swamp forest soil | GCA_024505145.1 |
| 63 | *Streptomyces violaceusniger* Tu 4113 | - | soil | GCA_000147815.3 |
| 64 | *Streptomyces antimycoticus* NBRC 100767 | - | soil | GCA_009936315.1 |
| 65 | *Streptomyces antimycoticus* NBRC 12839 | - | soil | GCA_005405925.1 |
| 66 | *Streptomyces* sp. AgN23 | - | rhizosphere | GCA_001598115.2 |
| 67 | *Streptomyces* sp. NRRL 30748 | - | soil | 39 |
| 68 | *Streptomyces* sp. M56 | - | termite-associated | 40 |
| 69 | *Streptomyces* sp. CWJ-256 | - | plant-associated | 41 |
| 70 | *Streptomyces* sp. 92JF-1 | - | marine | 42 |
| 71 | *Streptomyces* sp. KIB-H869 | - | plant-associated | 43 |
| 72 | *Streptomyces hygroscopicus* MSU-625 | - | soil | 44 |
| 73 | *Streptomyces hygroscopicus* MSU-616 | - | soil | 45 |
| 74 | *Streptomyces hygroscopicus* OUPS-N92 | - | marine | 46 |
| 75 | *Streptomyces* sp. CBR53 | - | - | 47 |
| 76 | *Streptomyces violaceusniger* NBRC 13459 | - | - | GCA_005405945.1 |
| 77 | *Streptomyces violaceusniger* NRRL F-8817 | - | - | GCA_001509775.1 |
| 78 | *Streptomyces* sp. HKI-0113 | - | - | 48 |
| 79 | *Streptomyces* sp. HKI-0114 | - | - | 48 |
| 80 | *Streptomyces* sp. 57-13 | - | - | 49 |
| 81 | *Streptomyces javensis* | - | - | GCA_016103505.1 |

"-": information missing.

**Tab. 5.** Summary of strains and plasmids used in this study.

| Strains | Description | Source/[Ref] |
|---|---|---|
| One Shot™ Mach1™ T1 Phage-Resistant Chemically Competent *E. coli* | For routine plasmids maintenance and cloning | Thermo Fisher Scientific |
| *E. coli* ET12567/pUZ8002 | For conjugating plasmids into *Streptomyces* | [50] |
| *S. iranensis* | Wild-type strain | DSMZ |
| *S. iranensis*/Δ*ptaA* | Δ*ptaA* mutant strain | In this work |
| *S. iranensis*/Δ*ptaA*/1J23 | Complementation strain of Δ*ptaA* mutant | In this work |
| *S. iranensis*/Δ*ptaA*/6M10 | Complementation strain of Δ*ptaA* mutant | In this work |
| *S. iranensis*/M2089I + E2090K + D2091M | TE domain mutant strain | In this work |
| *S. rapamycinicus* NRRL 5491 | Wild-type strain | DSMZ |
| *S. violaceusniger* Tu 4113 | Wild-type strain | DSMZ |
| *S. albus* DSM 41398 | Wild-type strain | DSMZ |
| **Plasmids** | | |
| pCRISPR-cBEST | For C to T base editing | [51] |
| pCRISPR-cBEST/Δ*ptaA* | Modified plasmid for inactivation of *ptaA* | In this work |
| pCRISPR-cBEST/ M2089I + E2090K + D2091M | Modified plasmid for site-specific mutation of TE domain | In this work |
| pESCA13/1J23 | BAC for complementation | In this work |
| pESCA13/6M10 | BAC for complementation | In this work |

**Tab. 6**. $^{1}$H (800 MHz) NMR data for **1–2** (in MeOD).

| position | $\Delta_H$ (J in Hz) | |
|---|---|---|
| | **1** | **2** |
| 1 | - | - |
| 2 | 5.90 (d, 15.4) | 5.97 (d, 15.1) |
| 3 | 7.33 (dd, 15.4, 11.1) | 7.16 (dd, 15.1, 10.9) |
| 4 | 6.26 (dd, 15.2, 10.8) | 6.25 (dd, 15.1, 10.9) |
| 5 | 6.13 (dd, 15.3, 8.8) | 6.07 (dd, 15.1, 8.6) |
| 6 | 2.48 (m) | 2.49 (m) |
| 7 | 3.85 (dd, 10.2, 2.2) | 3.32 (m) |
| 8 | 2.01 (m) | 2.02 (m) |
| 9 | 3.69 (dd, 11.4, 4.9) | 3.56 (dd, 11.5, 4.7) |
| 10 | 1.61 (m) | 1.69 (m) |
| 11 | - | - |
| 12 | 2.29 (dd, 14.9, 6.1), 1.64 (dd, 14.9, 1.9) | 2.19 (dd, 13.1, 4.3) 1.32 (dd, 13.2, 11.2) |
| 13 | 3.59 (m) | 3.66 (td, 10.8, 4.3) |
| 14 | 1.49 (m) | 1.02 (m) |
| 15 | 3.43 (m) | 3.88 (m) |
| 16 | 1.21 (d, 6.1) | 1.14 (d, 6.2) |
| 17 | 1.01 (d, 6.8) | 1.02 (d, 6.8) |
| 18 | 0.91 (d, 7.0) | 0.95 (d, 6.9) |
| 19 | 0.96 (d, 6.8) | 0.98 (d, 6.8) |
| 20 | 1.52 (m), 1.21 (m) | 1.60 (m), 1.44 (m) |
| 21 | 0.93 (t, 7.3) | 0.82 (t, 7.6) |

**Tab. 7**. $^{13}C$ (200 MHz) NMR data for **1–2** (in MeOD).

| position | $\Delta_C$, type | |
|---|---|---|
| | **1** | **2** |
| 1 | 168.7 | 170.2 |
| 2 | 120.6 | 122.9 |
| 3 | 146.7 | 148.1 |
| 4 | 129.6 | 129.6 |
| 5 | 151.0 | 148.1 |
| 6 | 40.4 | 40.5 |
| 7 | 75.5 | 78.0 |
| 8 | 37.5 | 38.0 |
| 9 | 72.8 | 74.7 |
| 10 | 42.2 | 42.1 |
| 11 | 103.2 | 103.2 |
| 12 | 37.4 | 33.8 |
| 13 | 70.3 | 66.3 |
| 14 | 51.3 | 52.2 |
| 15 | 72.7 | 66.3 |
| 16 | 20.9 | 20.4 |
| 17 | 16.0 | 15.9 |
| 18 | 5.0 | 5.3 |
| 19 | 12.1 | 12.7 |
| 20 | 24.8 | 19.7 |
| 21 | 10.1 | 10.5 |

**Tab. 8.** Summary of primers used in this study.

| Primer name | Sequence (5' → 3') | Description |
|---|---|---|
| SOS1-F | TTACCAGCCCCCAAGAAACG | Forward primer for qRT-PCR to detect the relative expression of SOS1 |
| SOS1-R | TCAACTGTAGGCCAGTCAGC | Reverse primer for qRT-PCR to detect the relative expression of SOS1 |
| TIP2;3-F | TAATGGCAAGAGCGTACCGAC | Forward primer for qRT-PCR to detect the relative expression of TIP2;3 |
| TIP2;3-R | ACCAATGCAAAGGTCACAACG | Reverse primer for qRT-PCR to detect the relative expression of TIP2;3 |
| Del-*ptaA* | CGGTTGGTAGGATCGACGGCGCACCCAGGCGGTATGCGTAGTTTTAGAGCTAGAAATAGC | Inactivation of *ptaA*, the base marked in red is sgRNA sequence |
| Mut-*ptaE* | CCGGTTGGTAGGATCGACGGGGTCCTCCATCATGGTGAAGGTTTTAGAGCTAGAAATAGC | Site-directed mutagenesis of TE domain in *ptaE*, the base marked in red is sgRNA sequence |
| ID-sgRNA-F | TGTGTGGAATTGTGAGCGGATA | Forward primer for screening plasmid |
| ID-sgRNA-R | CCCATTCAAGAACAGCAAGCA | Reverse primer for screening plasmid |
| ID-*ptaA*-F | TTGCACAGCTCGACGGACAT | Forward primer for screening *ptaA* mutants |
| ID-*ptaA*-R | GTGTCACCCGCTTTGTCGA | Reverse primer for screening *ptaA* mutants |
| ID-*ptaE*-F | CAACGCCATGATCGTCGTTC | Forward primer for screening TE domain mutants |
| ID-*ptaE*-R | CGTTCGAGACCGGGAAATG | Reverse primer for screening TE domain mutants |
| ID-1J23-right-F | GTCGACATGGCTTGCCTC | Forward primer for validating the right flank of *S. iranensis*/Δ*ptaA*/1J23 |
| ID-1J23-right-R | ATCCGTCTCGACTCCGG | Reverse primer for validating the right flank of *S. iranensis*/Δ*ptaA*/1J23 |
| ID-1J23-left-F | AGCAGAAGGTAGGGCAG | Forward primer for validating the left flank of *S. iranensis*/Δ*ptaA*/1J23 |
| ID-1J23-left-R | GAGGAGACTTCTGCCATGTC | Reverse primer for validating the left flank of *S. iranensis*/Δ*ptaA*/1J23 |
| ID-6M10-right-F | GATCTGCTGCTGTTCACGG | Forward primer for validating the right flank of *S. iranensis*/Δ*ptaA*/6M10 |
| ID-6M10-right-R | CCGAGCAGATCCGAGATG | Reverse primer for validating the right flank of *S. iranensis*/Δ*ptaA*/6M10 |
| ID-6M10-left-F | GAGCACCATCAGCAGGCG | Forward primer for validating the left flank of *S. iranensis*/Δ*ptaA*/6M10 |
| ID-6M10-left-R | CATGATGTCCGTGTCGCTC | Reverse primer for validating the left flank of *S. iranensis*/Δ*ptaA*/6M10 |

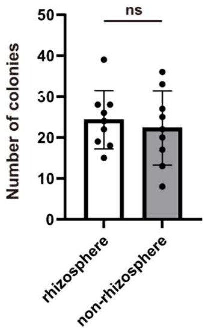


**Fig. 1**. Enrichment evaluation of *S. iranensis* in rhizosphere soil (mean ± SD, n=9). Statistical significance was assessed by unpaired t test.

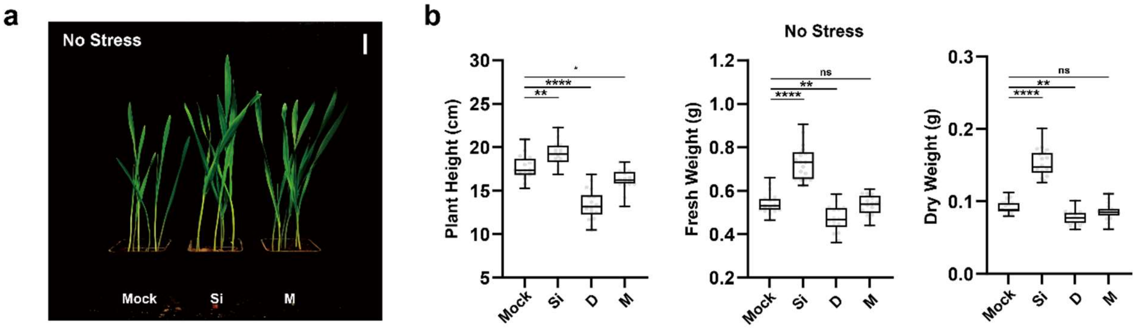


**Fig. 2**. Barley experiments of *S. iranensis* and *S. iranensis*/Δ*ptaA* treatment under non-stress condition. ***a***, *S. iranensis* and its plant growth-promoting activity on barley seedlings growth (bars=2 cm); ***b***, the box-plots (middle bar=median, box limit=upper and lower quartile, extremes=min and max values) depict the plant height, fresh weight and dry weight of barley seedlings growing on non-stress condition (n =18); Abbreviation: ***Mock***, control; ***Si***, treatment of *S. iranensis* culture broth; ***D***, treatment of *S. iranensis*/Δ*ptaA* culture broth; ***M***, treatment of blank medium (ISP2). Statistical significance was assessed by one-way ANOVA with post hoc Dunnett's multiple comparisons test. Asterisks indicate the level of statistical significance: $^{*}P < 0.05$, $^{**}P < 0.01$, $^{***}P < 0.001$ and $^{****}P < 0.0001$.

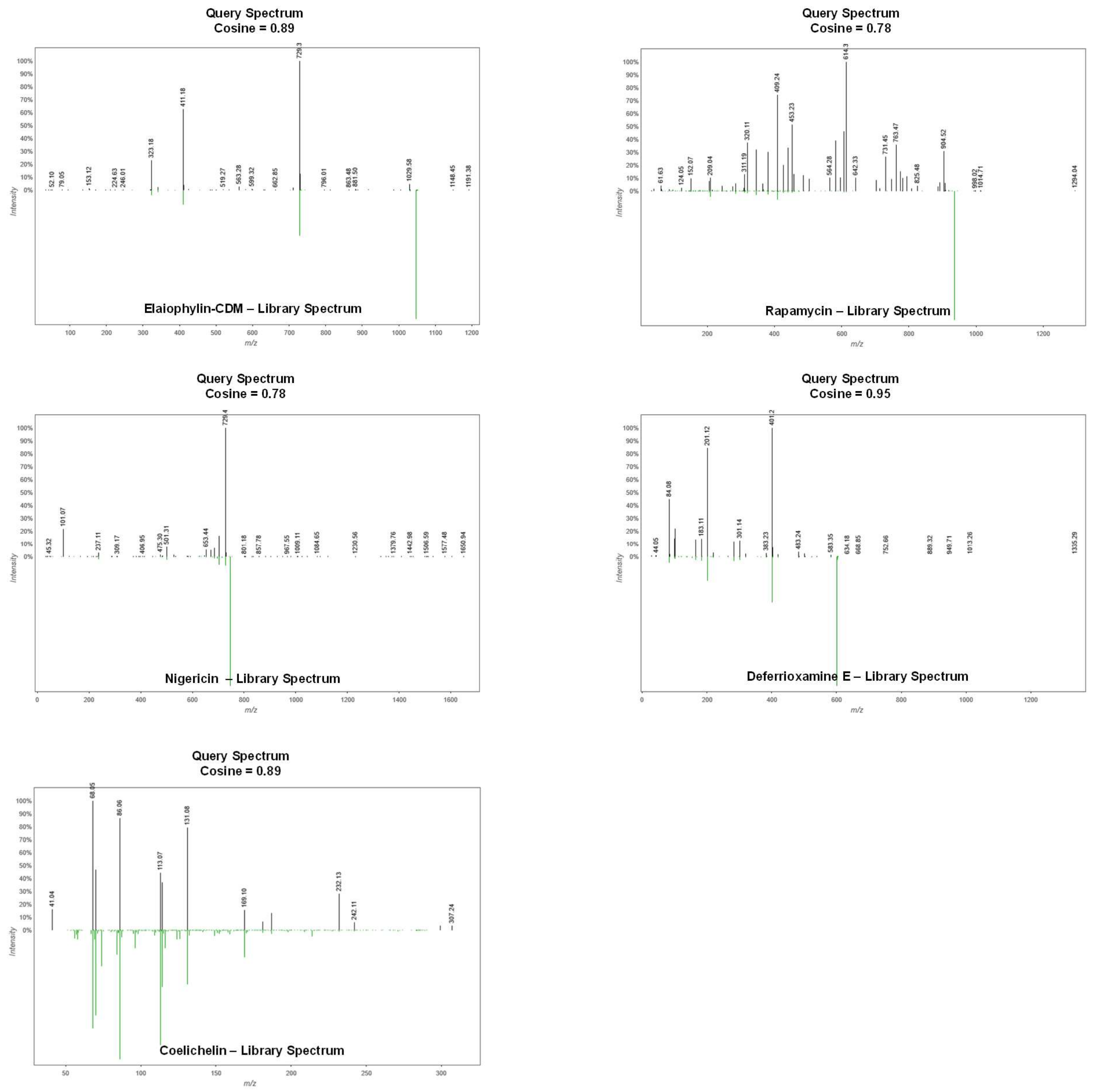


**Fig. 3**. Mirror plots comparing spectra from known metabolites from *S. iranensis* to standard spectra deposited in GNPS. In the upper part of the plot (black lines) is represented the MS spectra of the candidate feature and in the lower part (green lines) is the MS spectra of the standard compound. Mirror plots have been generated using https://metabolomics-usi.ucsd.edu/.

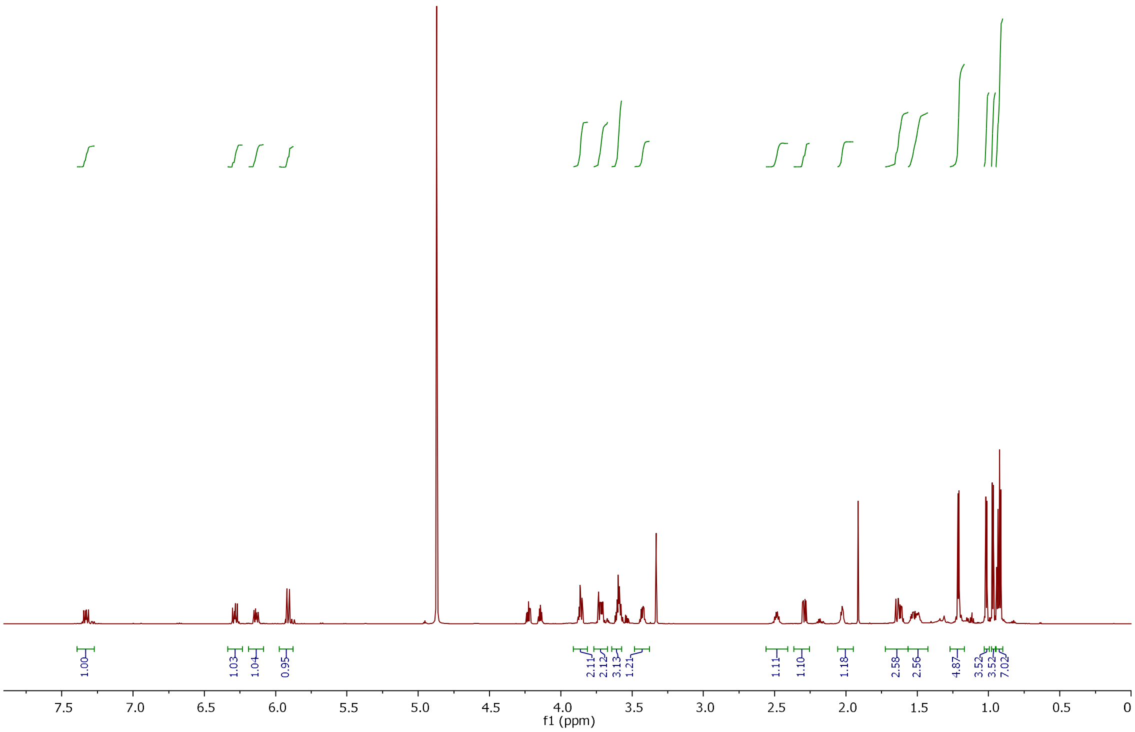

**Fig. 4**. $^{1}$H NMR spectrum of **1.**

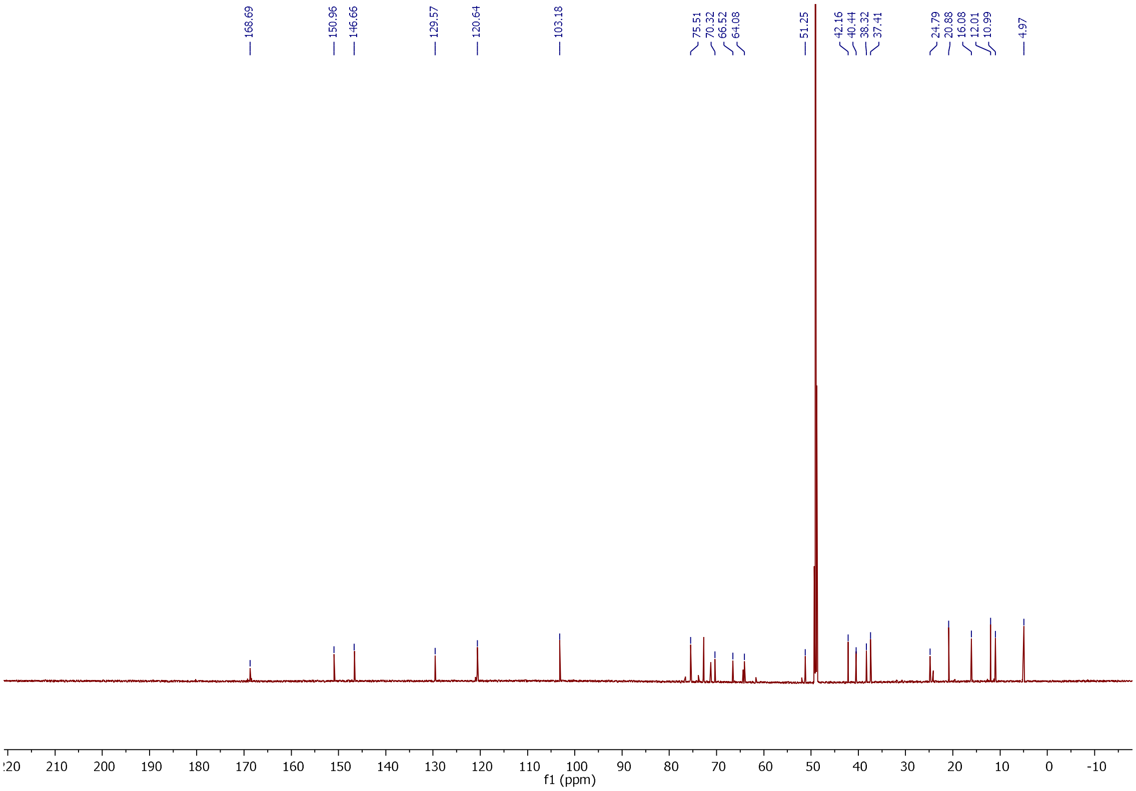

**Fig. 5**. $^{13}$C NMR spectrum of **1.**

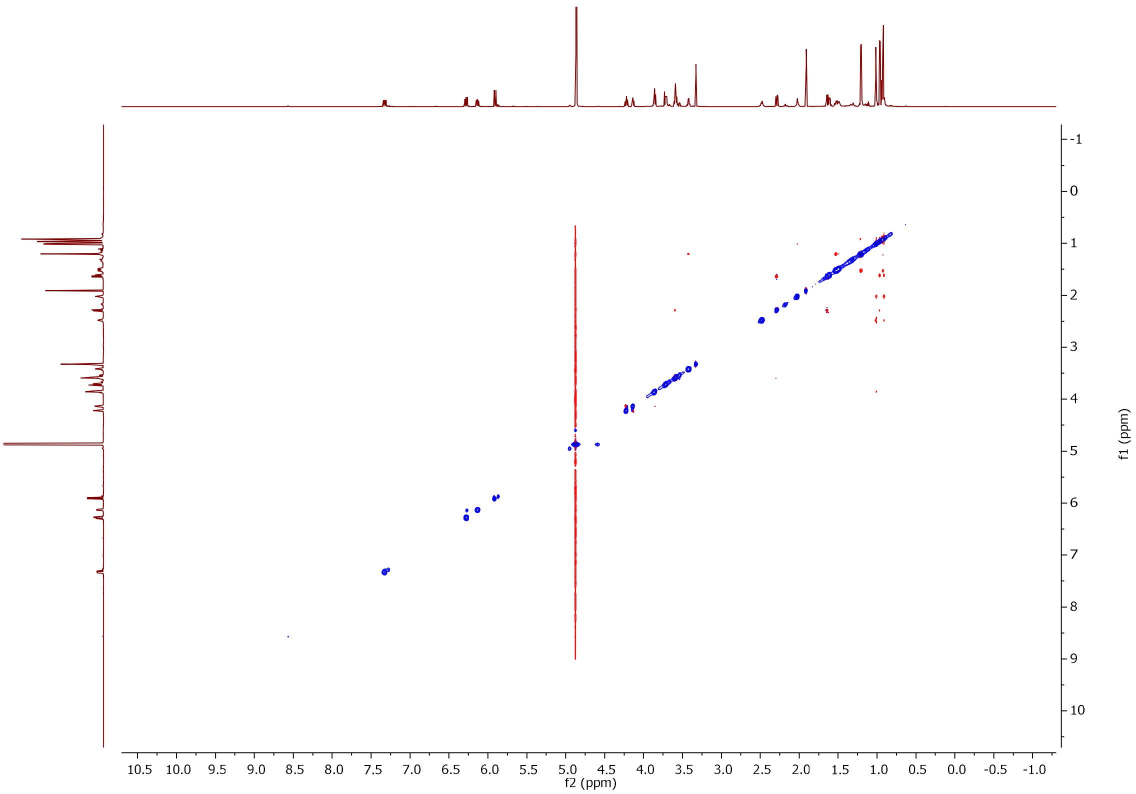


**Fig. 6**. NOESY spectrum of **1**.

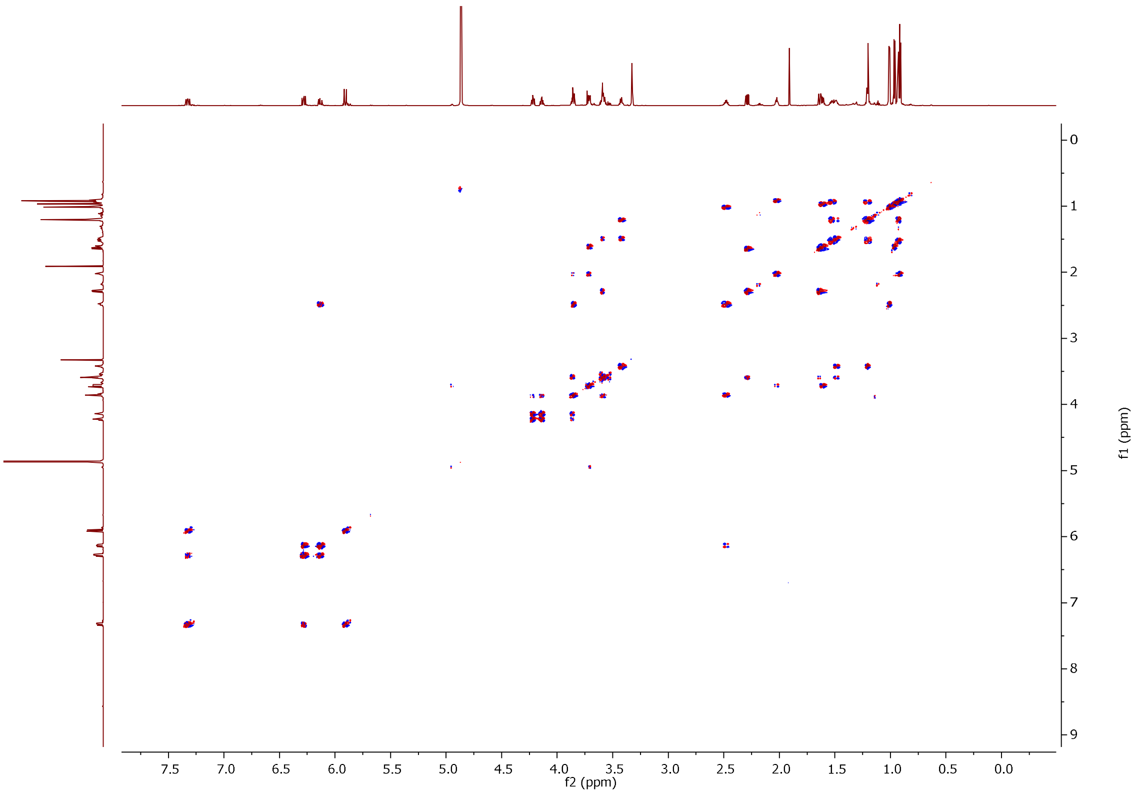


**Fig. 7.** COSY spectrum of **1**.

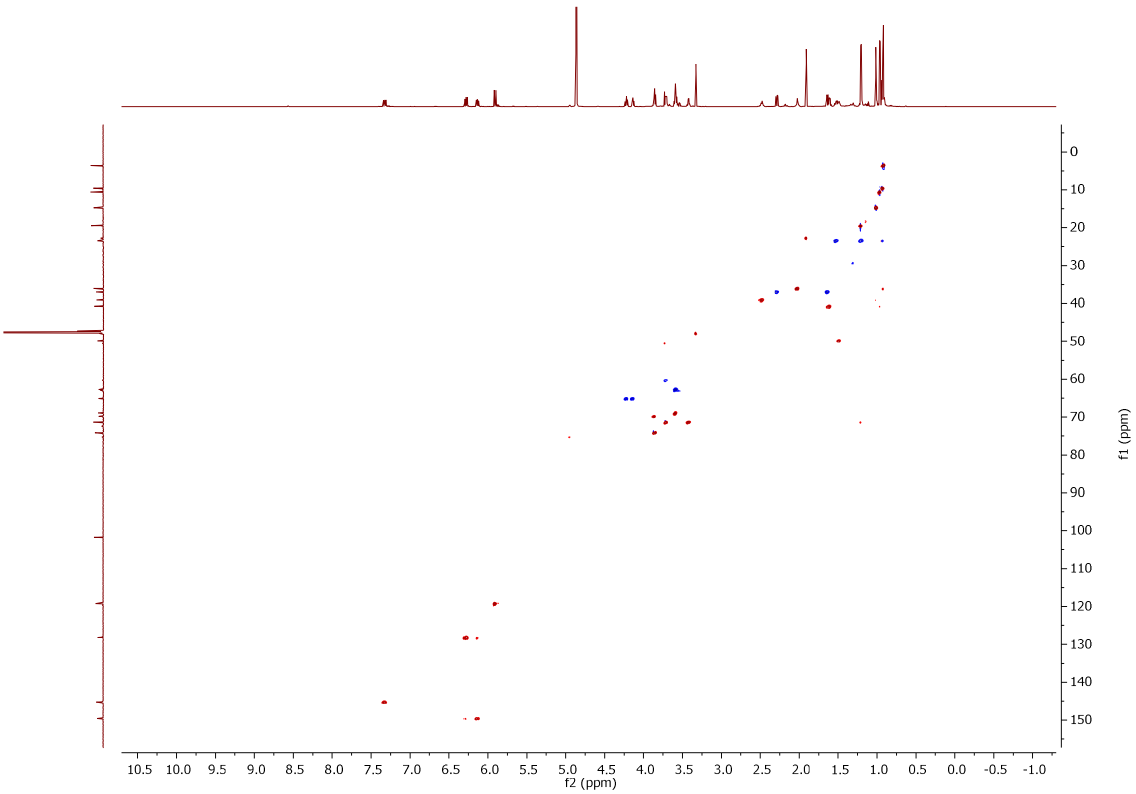


**Fig. 8.** HSQC spectrum of **1**.

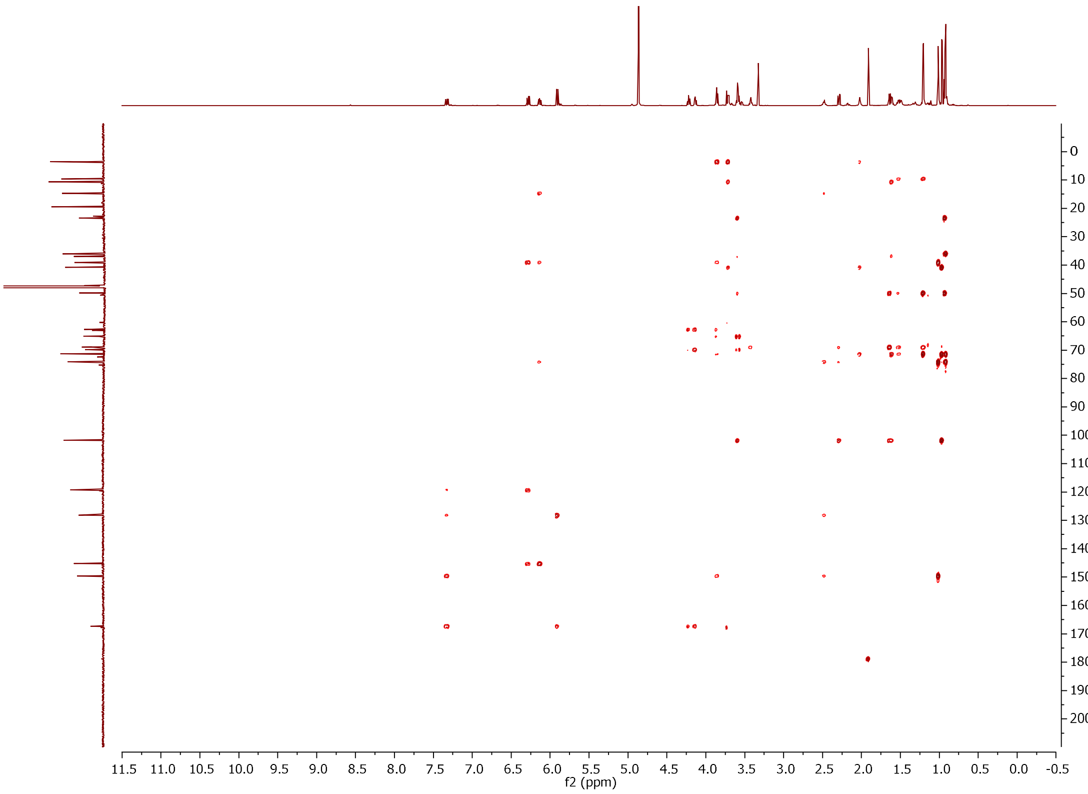


**Fig. 9**. HMBC spectrum of **1**.

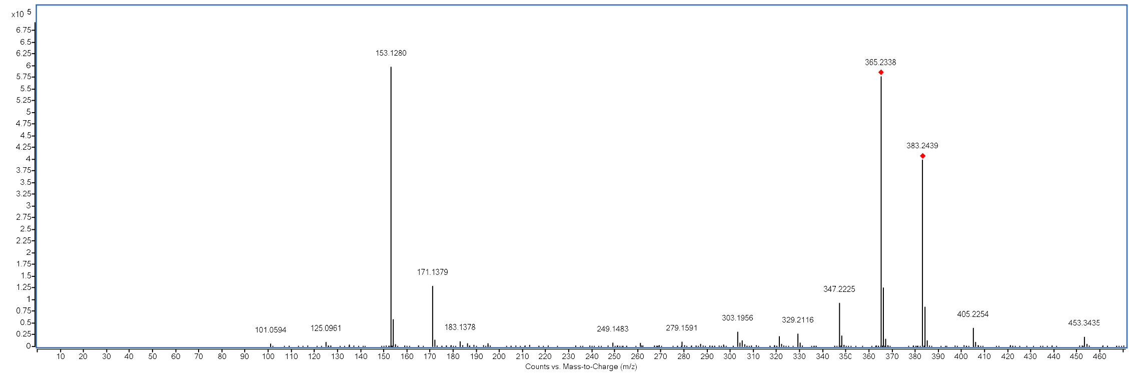


**Fig. 10**. Mass spectrum of **1**. Its formula of $C_{21}H_{34}O_6$ was deduced by *m/z* 383.2439 $[M+H]^+$ (calculated for 383.2428, *Δ* 2.84 ppm).

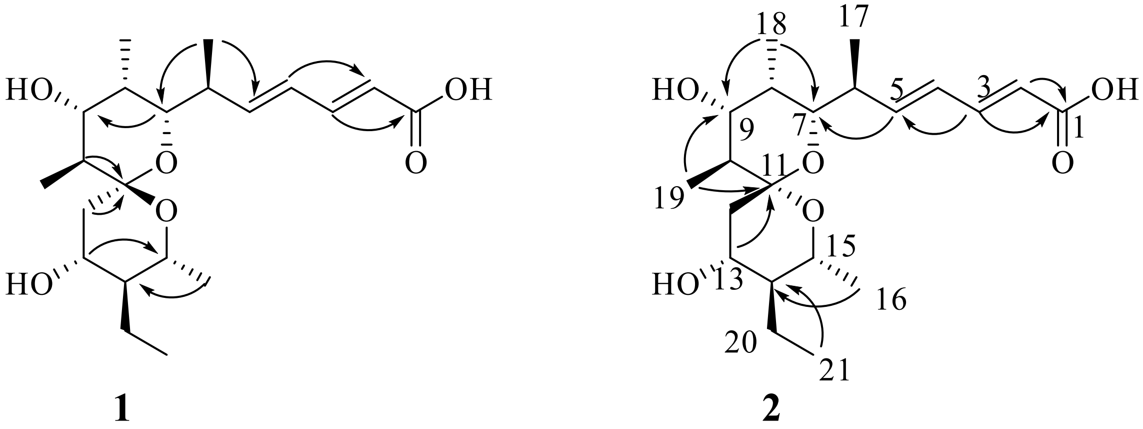


**Fig. 11**. Selected HMBC correlations for **1** and **2**.

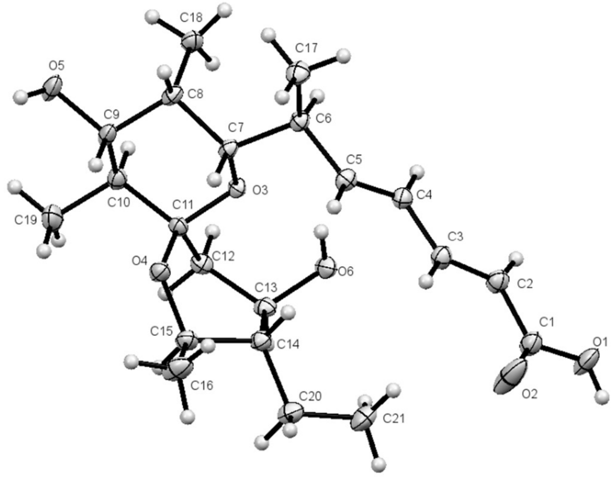


**Fig. 12**. Crystal structure of **1**. ORTEP diagram showing the atom-numbering scheme and solid-state conformation of **1**.

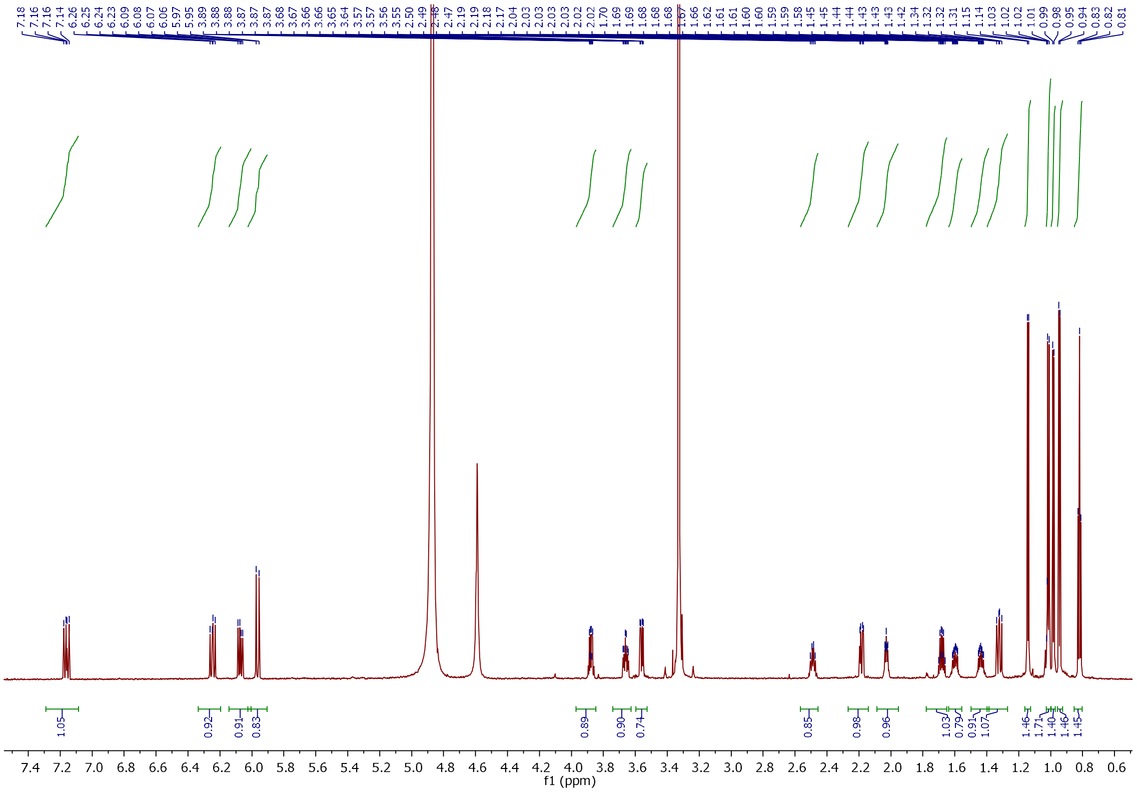

Fig. 13. $^{1}$H NMR spectrum of **2**.

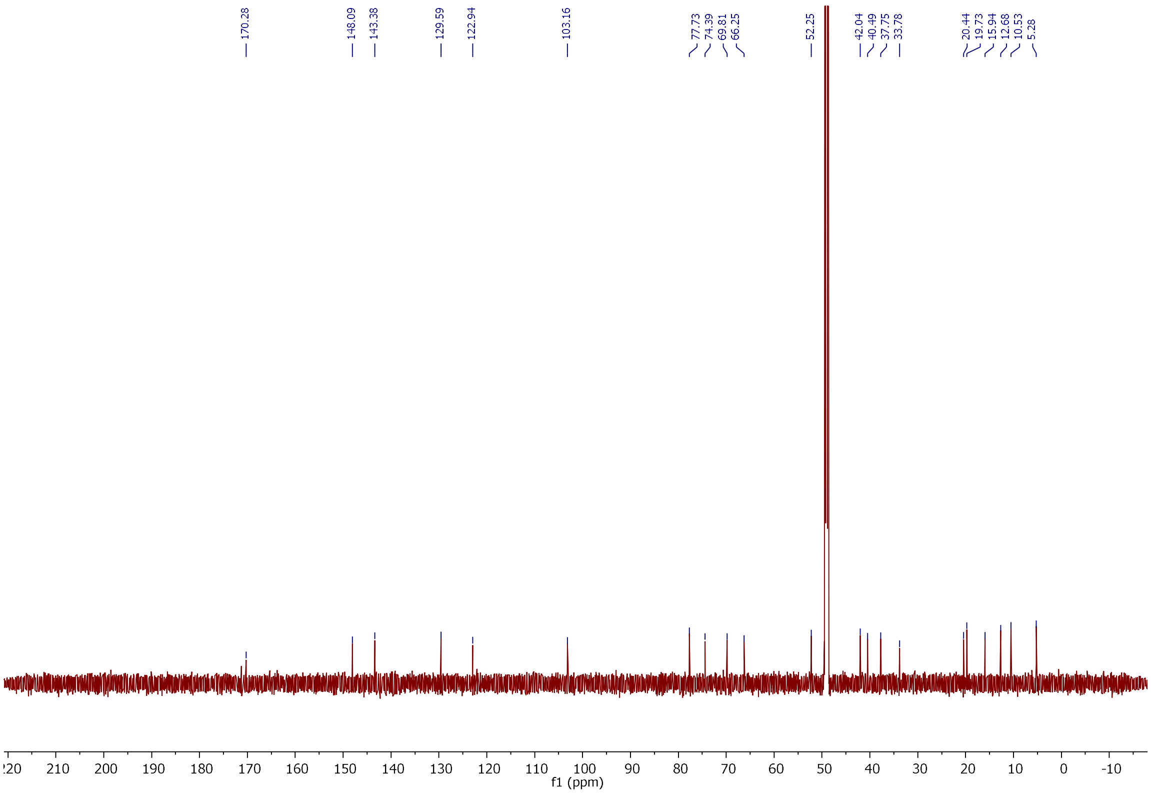

Fig. 14. $^{13}$C NMR spectrum of **2**.

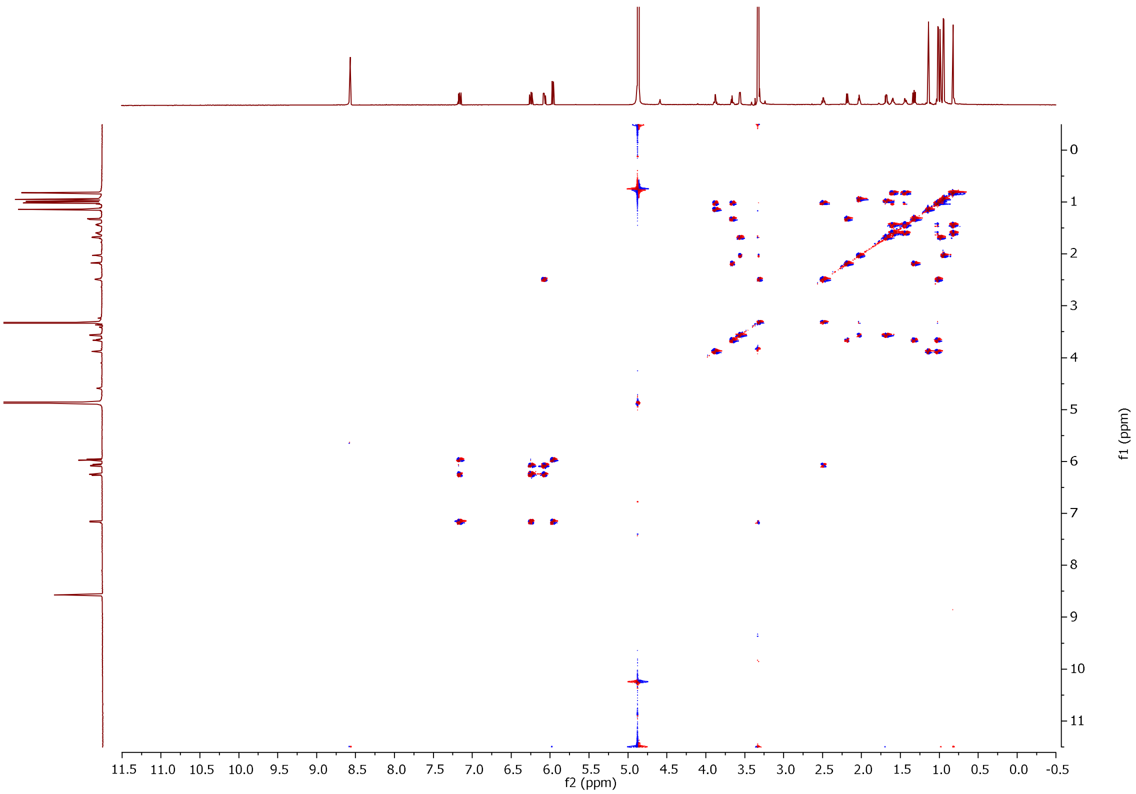


**Fig. 15.** COSY spectrum of **2**.

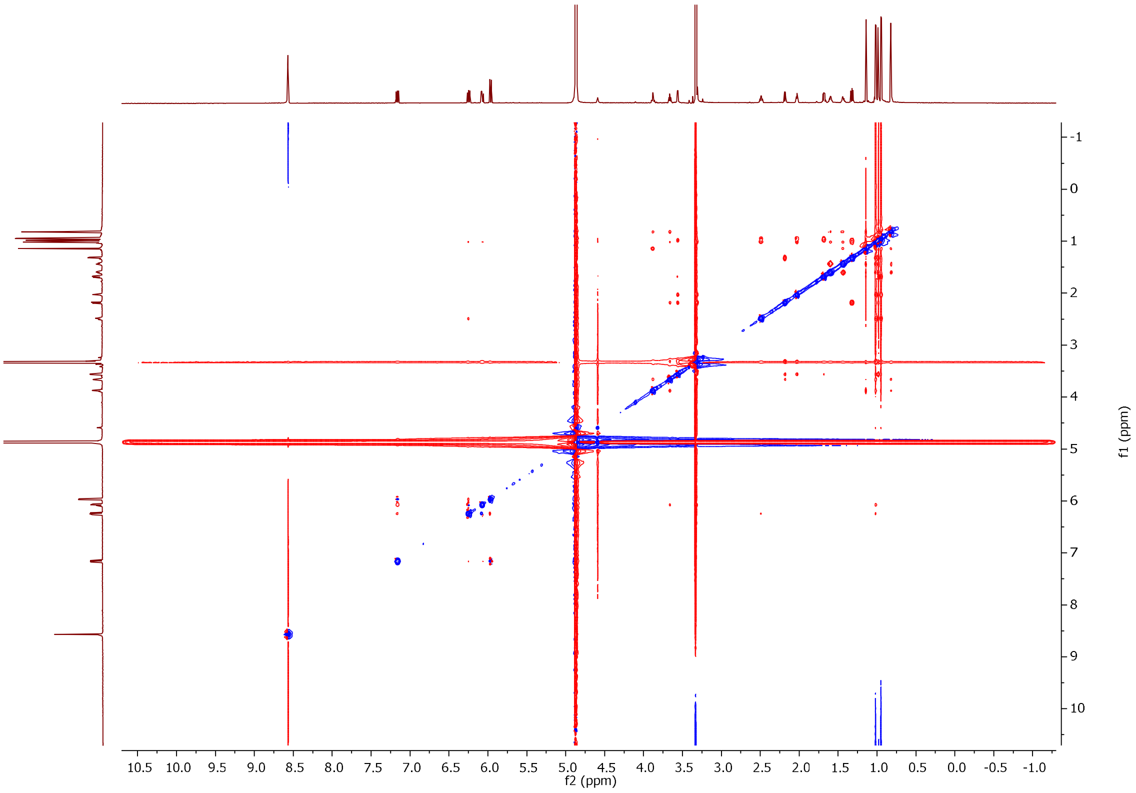


**Fig. 16.** NOESY spectrum of **2**.

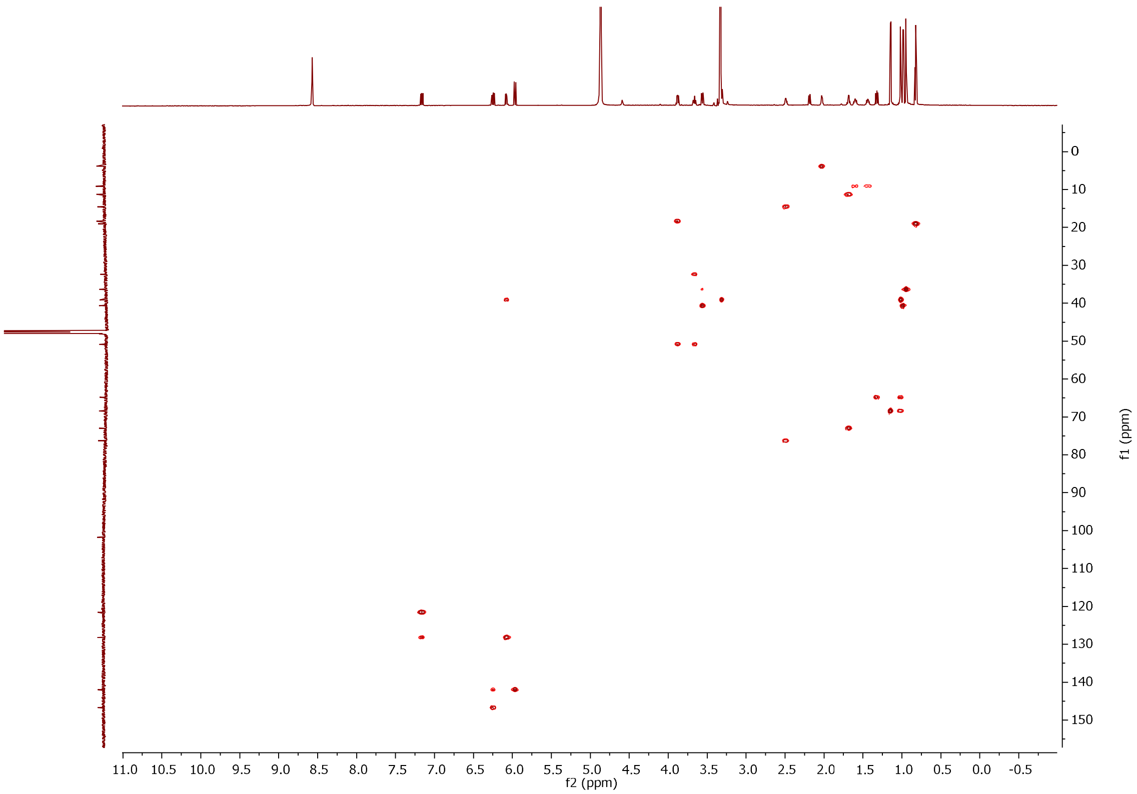


**Fig. 17**. H2BC spectrum of compound **2**.

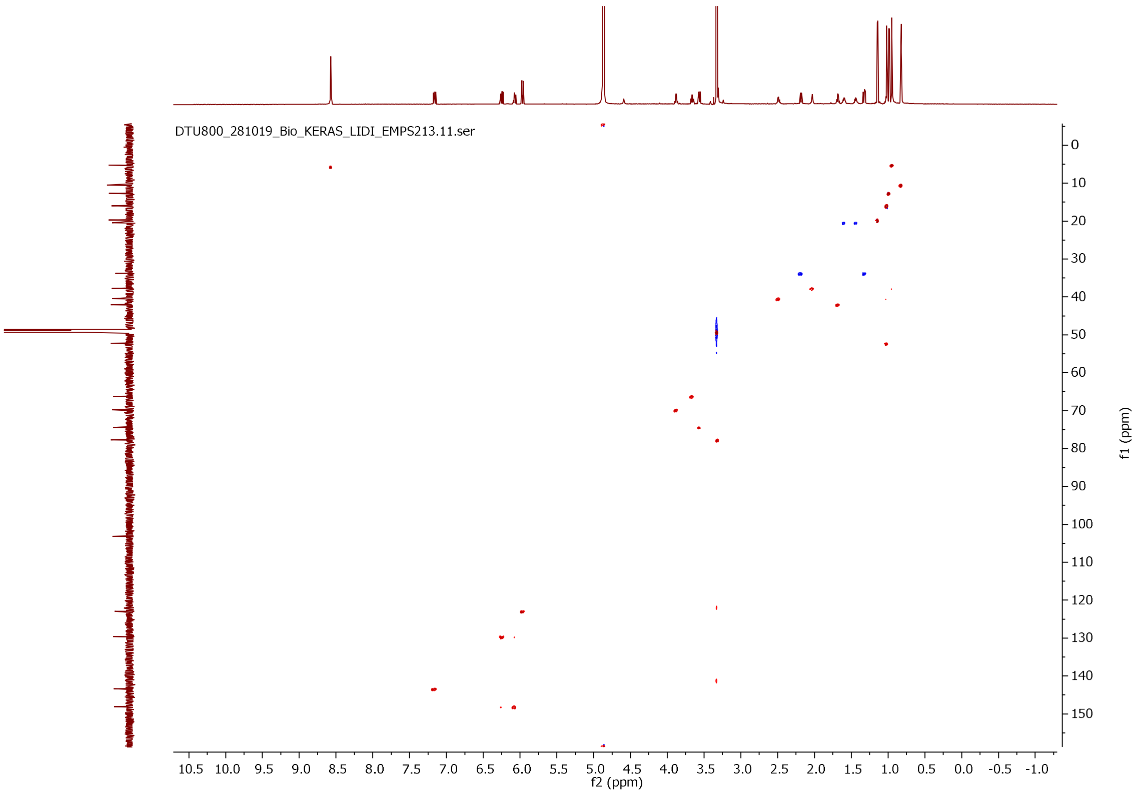


**Fig. 18.** HSQC spectrum of **2**.

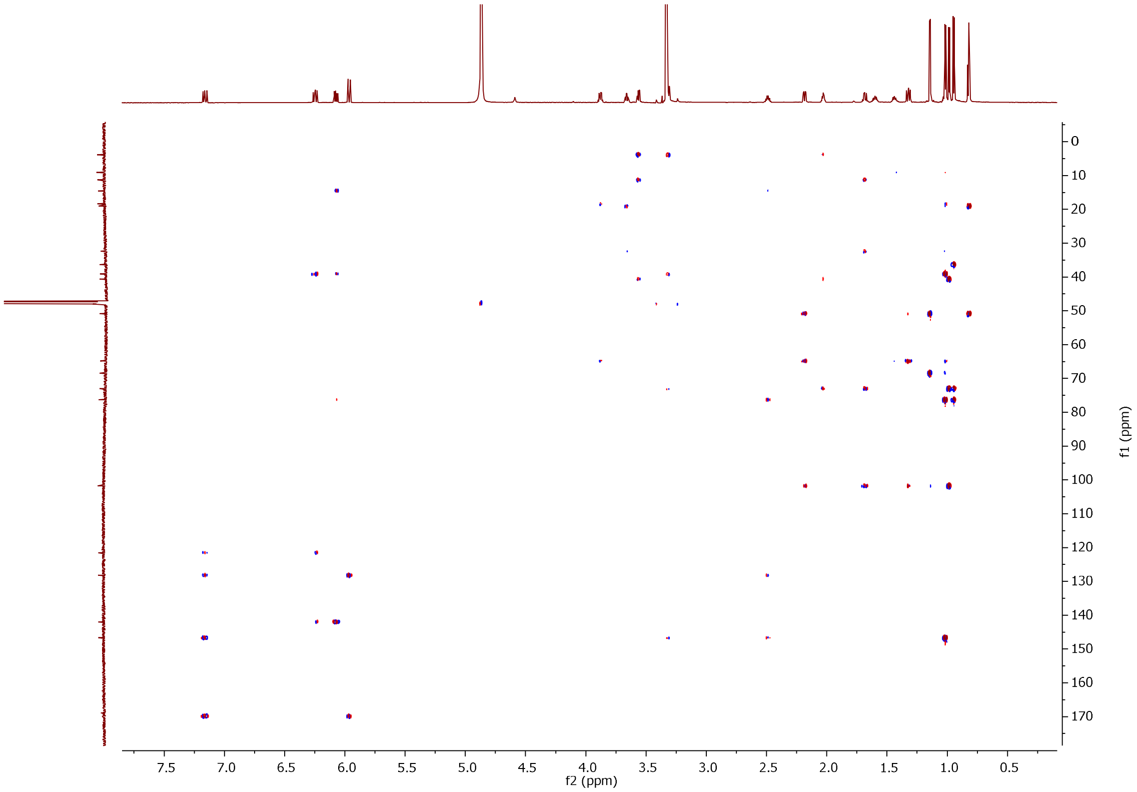


**Fig. 19.** HMBC spectrum of **2**.

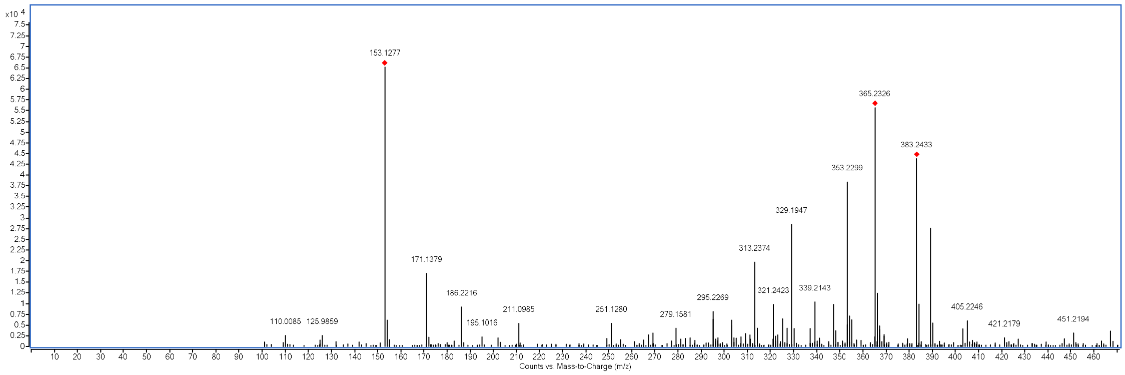


**Fig. 20**. Mass spectrum of **2**. Its formula of $C_{21}H_{34}O_6$ was deduced by *m/z* 383.2433 $[M+H]^+$ (calculated for 383.2428, *Δ* 1.27 ppm).

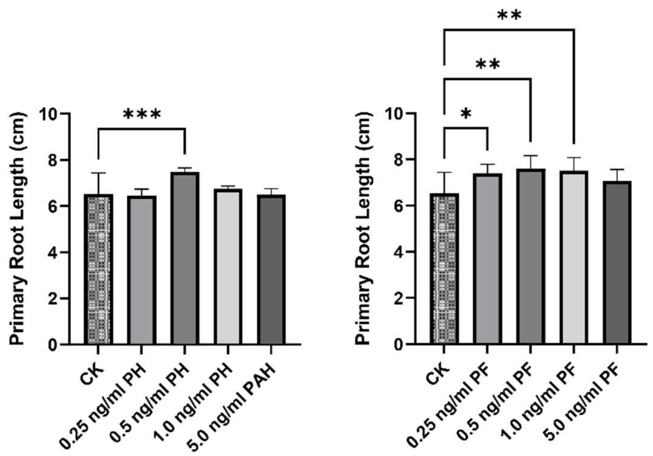


**Fig. 21**. Primary root length of *Arabidopsis* seedling treated with different concentrations of pteridic acids H and F. Abbreviation: CK, blank control treated by sterile Milli-Q water; PH, treatment with pteridic acid H (mean ± SD, n=8). Statistical significance was assessed by one-way ANOVA with post hoc Dunnett's multiple comparisons test. Asterisks indicate the level of statistical significance: *$p < 0.05$, **$p < 0.01$, ***$p < 0.001$.

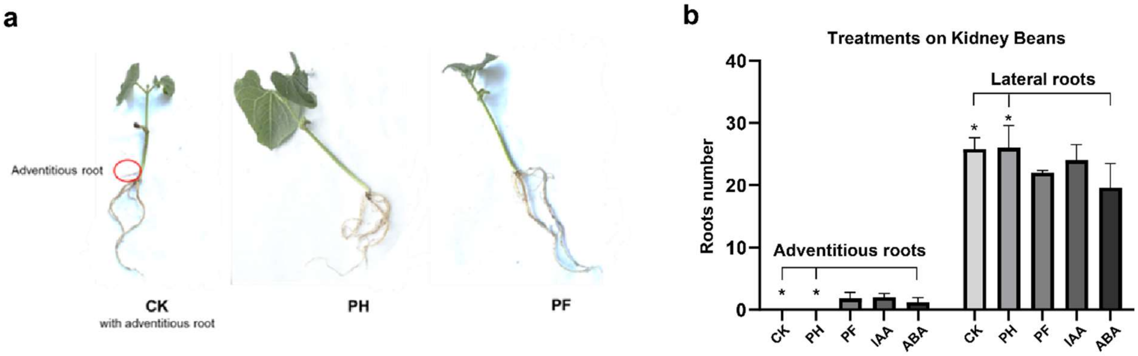


**Fig. 22**. Kidney beans growth experiment with pure pteridic acids. ***a***, the phenotypes of Kidney beans after treatments with pteridic acids H and F. ***b***, the numbers of adventitious roots and lateral roots of Kidney beans after different treatments (mean ± SD, n=5). Abbreviation: ***CK***, control; ***PH***, treatment of 1 ng $mL^{-1}$ pteridic acid H; ***PF***, treatment of 1 ng $mL^{-1}$ pteridic acid F; ***IAA***, treatment of 1 ng $mL^{-1}$ IAA; ***ABA***, treatment of 1 ng $mL^{-1}$ ABA. Asterisks indicate the level of statistical significance: *$P < 0.05$. Statistical significance was assessed by T-test.

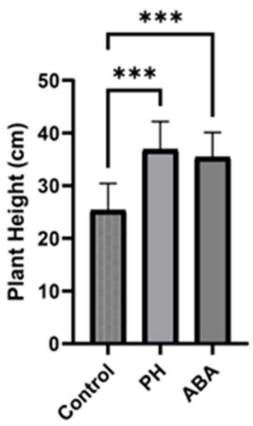


**Fig. 23**. Pteridic acid H and ABA at 1 ng $mL^{-1}$ help Mung beans against heavy metal stress. The plant height of Mung beans after treatments with pteridic acids H and ABA. Abbreviation: ***CK***, Control; ***PH***, treatment of pure pteridic acid H; ***ABA***, treatment of abscisic acid (mean ± SD, n=9). Asterisks indicate the level of statistical significance: ***$P < 0.001$. Statistical significance was assessed by one-way ANOVA with post hoc Dunnett's multiple comparisons test.

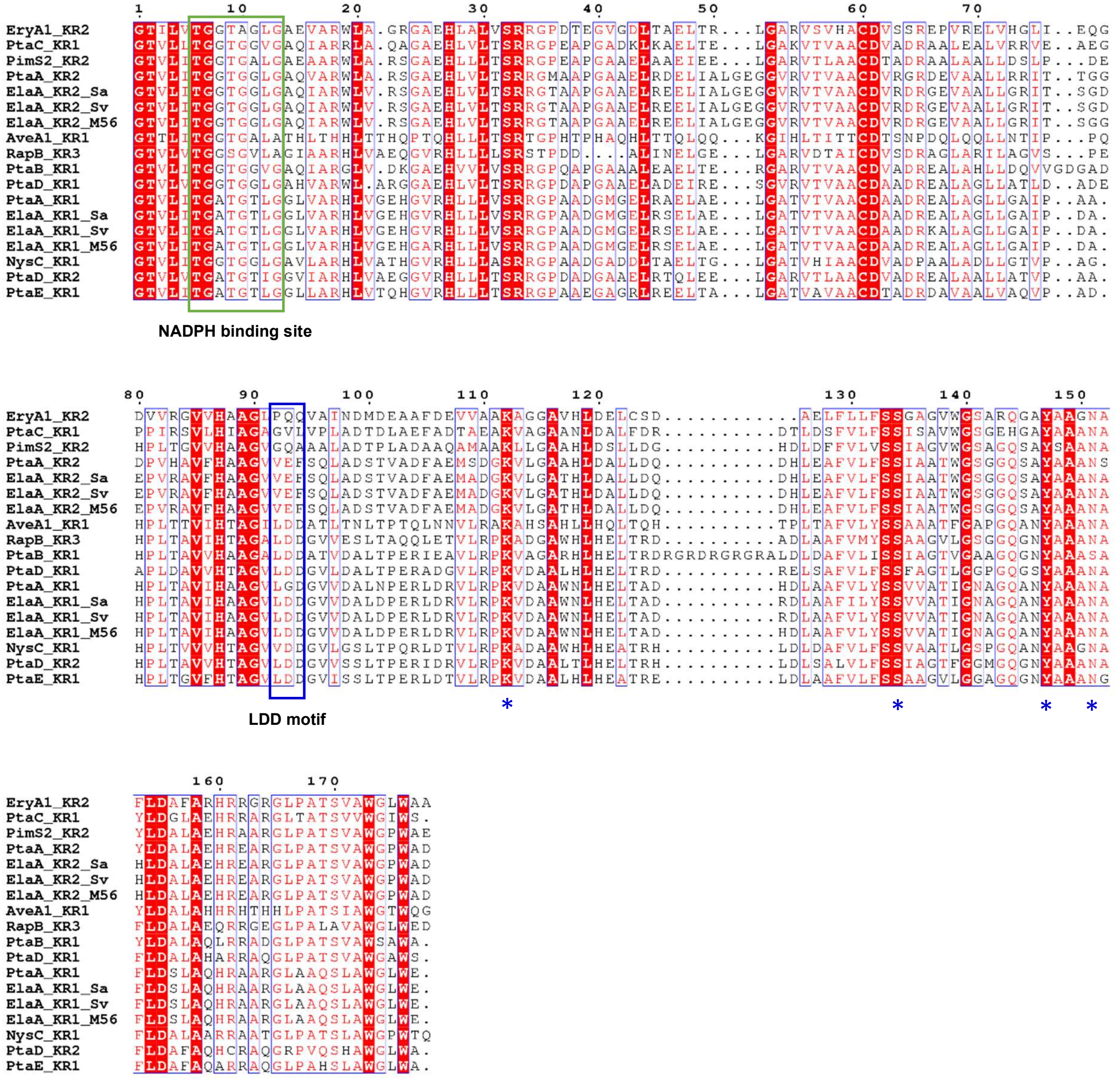


**Fig. 24**. Multiple sequence alignment of KR domains. Green box indicates NADPH binding site and blue box indicates the LDD motif. Abbreviation: Ery, erythromycin; Pta, pteridic acids; Pim, pimaricin; Ave, avermectin; Rap, rapamycin; Nys, nystatin.

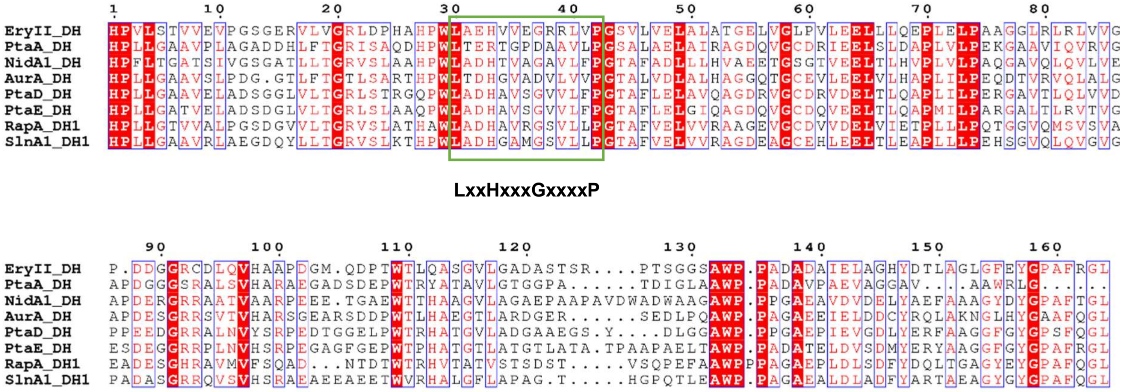

**Fig. 25**. Multiple sequence alignment of DH domains. The green box indicates the conserved LxxHxxGxxxxP motif. Abbreviation: Ery, erythromycin; Pta, pteridic acids; Nid, niddamycin; Aur, aureothin; Rap, rapamycin; Sln, salinomycin.

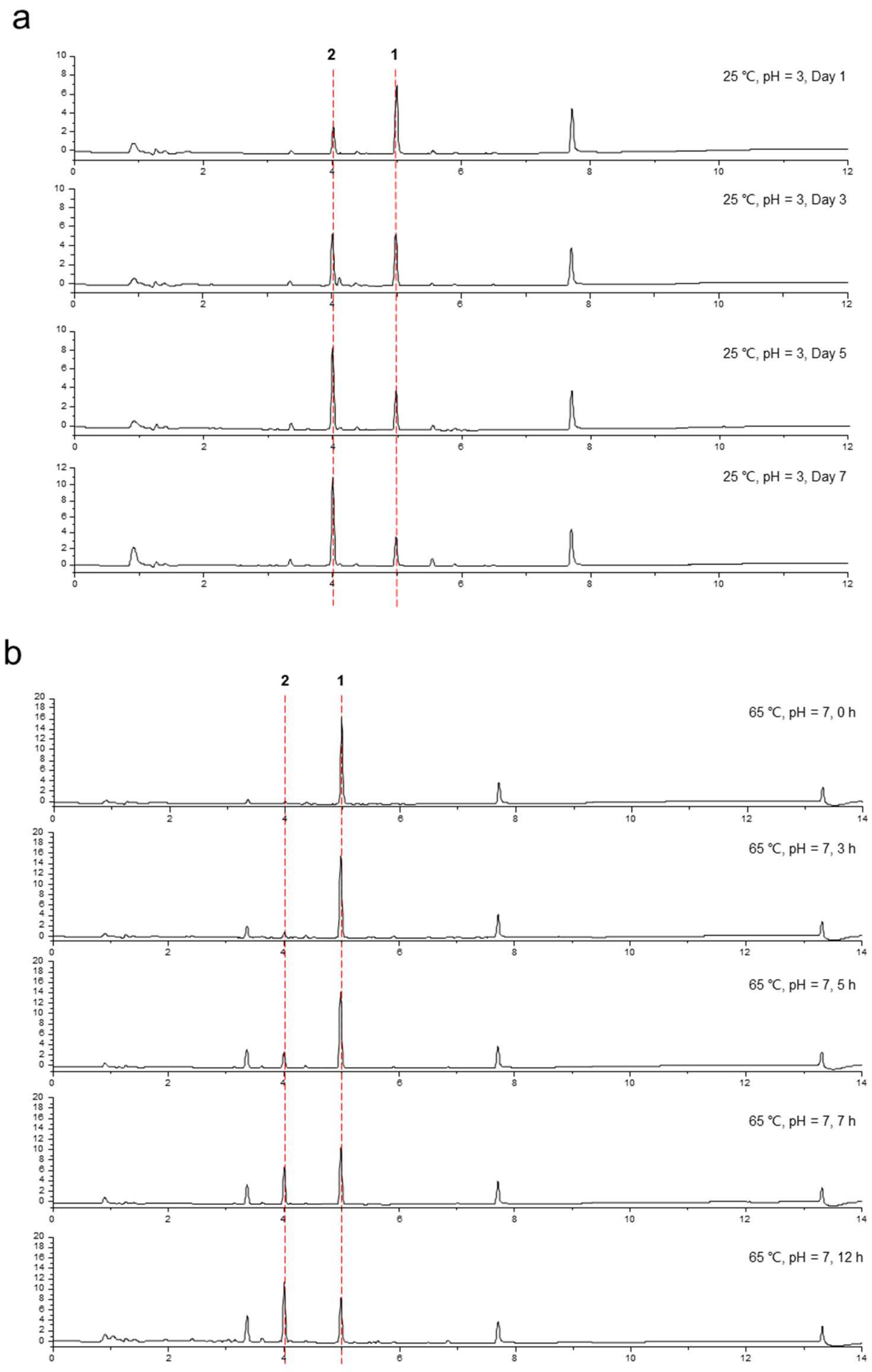


**Fig. 26**. Stability test of pteridic acids. Abbreviation: **1**, pteridic acid H; **2**, pteridic acid F. **1** was tested unstable in pH 3 buffer solution at 25 ˚C. ***a***, **1** was transformed to **2** fast, after 3 days the contents of them were as equal. The transformation rate is approximately 12.5%, 25%, 37.5%, 50% in 1 d, 3 d, 5 d and 7 d in pH 3 buffer solution at 25 ˚C. ***b***, **1** was unstable in water with 65 ˚C and transformed to **2** fast, after 12 hours, the content of **1** exceeded **2**. The transformation rate is approximately 7 %, 33 % and 47 % in 3 h, 7 h and 12 h, 65 ˚C.

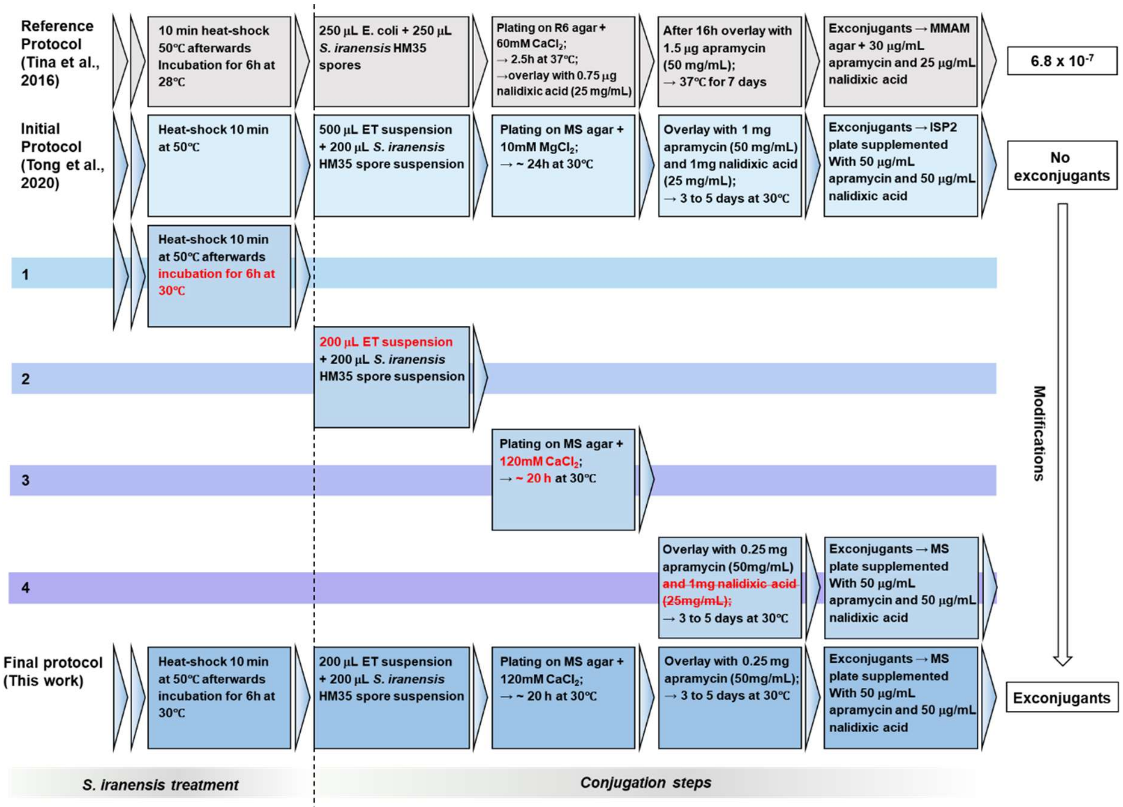


**Fig. 27**. The schematic of optimized genetic manipulation in *S. iranensis* by using CRISPR-cBEST system.

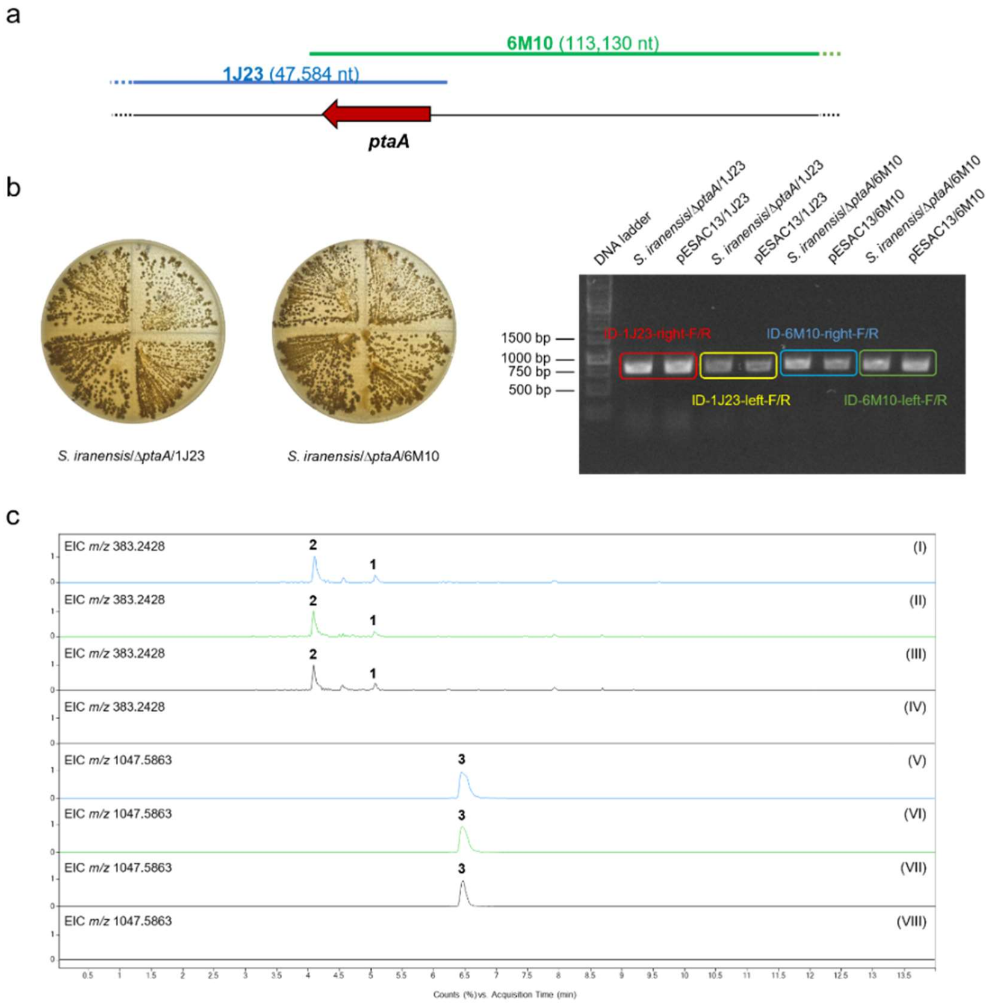


**Fig. 28**. Complementation experiment of *ptaA*-inactivation mutant of *S. iranensis*. (a) location of pESCA13/1J23 and pESCA13/6M10 in genome of *S.iranensis*. (b) apramycin-resistance screening and PCR verification of *S. iranensis*/Δ*ptaA*/1J23 and *S. iranensis*/Δ*ptaA*/6M10. (c) Extract Ion Chromatography (EIC) in positive mode was performed to detect pteridic acid H (1) and pteridic acid F (2) (*m/z* 383.2428 $[M+H]^+$ Δ ±5 ppm) as well as elaiophylin (3) (*m/z* 1047.5863 $[M+Na]^+$ Δ ±5 ppm) in the *S. iranensis*/Δ*ptaA*/1J23 (trace I and V), *S. iranensis*/Δ*ptaA*/6M10 (trace II and VI), wild-type *S. iranensis* (trace III and VII), and *S. iranensis*/Δ*ptaA* (trace IV and VIII).

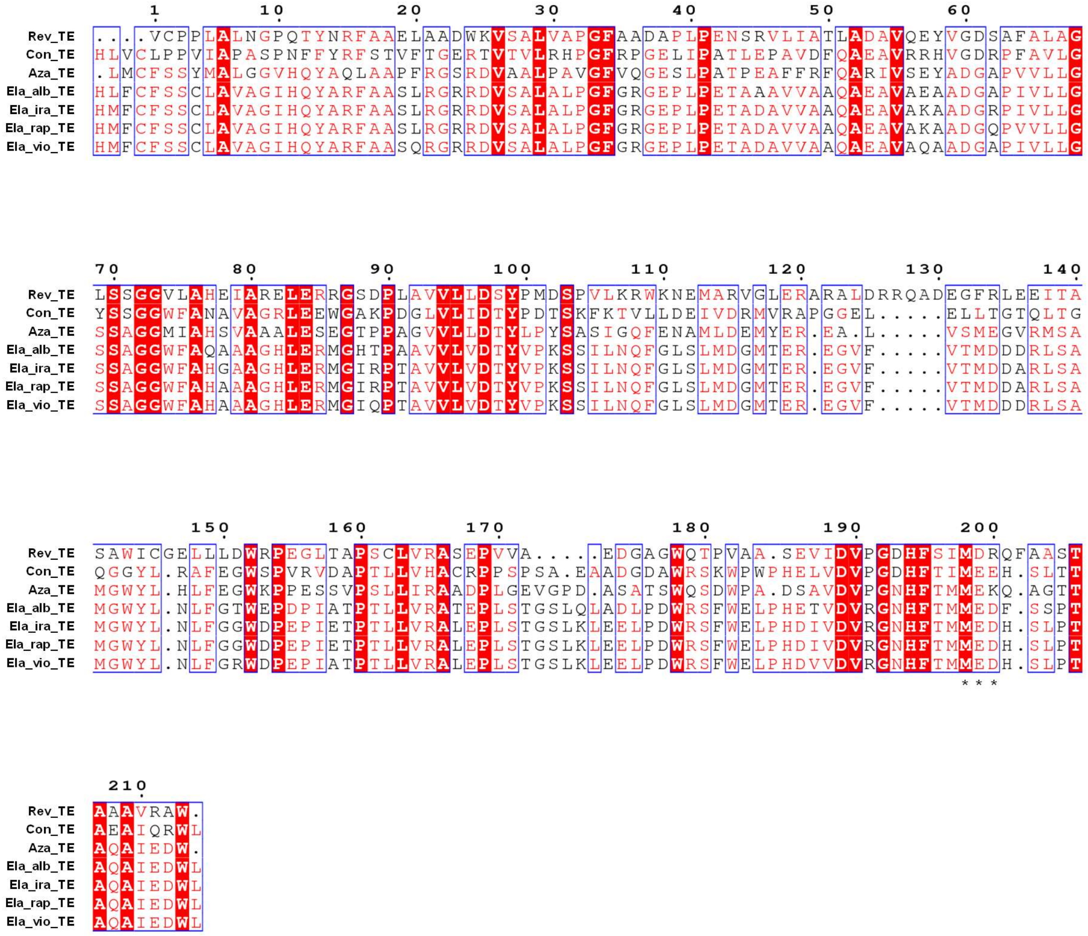


**Fig. 29**. Multiple sequence alignment of TE domains. The three amino acids marked with an asterisk were mutated in this study. Abbreviation: Rev_TE, the TE domain in reveromycin A biosynthesis from *Streptomyces* sp. SN-593; Con_TE, the TE domain in conglobatin biosynthesis from *Streptomyces conglobatus*; Aza_TE, the TE domain in azalomycin F3a biosynthesis from *Streptomyces* sp. 211726; Ela_alb_TE, the TE domain in elaiophylin biosynthesis from *S. albus* DSM 41398; Ela_ira_TE, the TE domain in elaiophylin biosynthesis from *S. iranensis* HM 35; Ela_rap_TE, the TE domain in elaiophylin biosynthesis from *S. rapamycinicus* NRRL 5491; Ela_vio_TE, the TE domain in elaiophylin biosynthesis from *S. violaceusniger* Tu 4113.

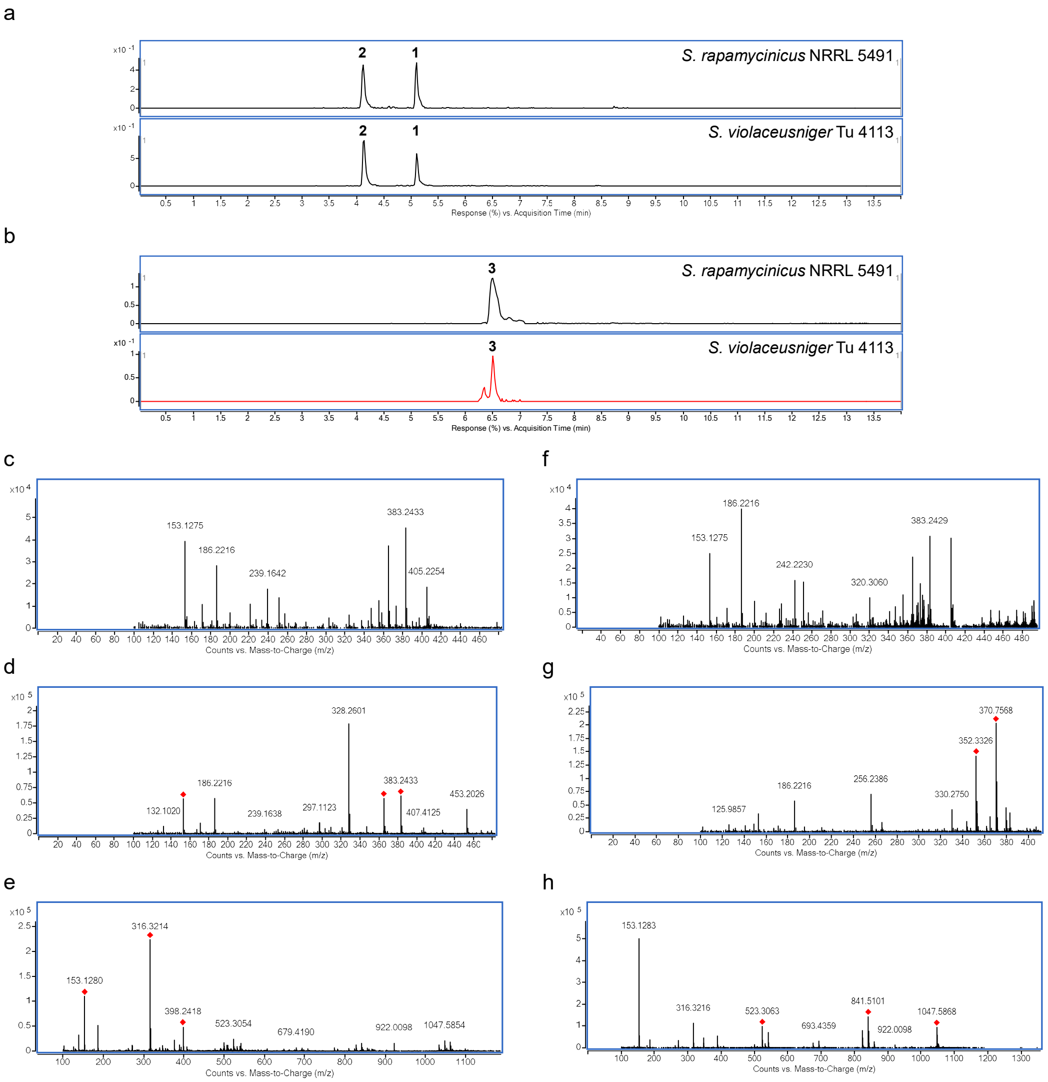


**Fig. 30**. The HR-LC-MS/MS analysis of metabolites in *S. rapamycinicus* NRRL 5491 and *S. violaceusniger* Tu 4113. ***a***, the Extracted Ion Chromatography (EIC) of *m/z* 383.2428 $[M+H]^+$ in wild-type *S. rapamycinicus* NRRL 5491 and *S. violaceusniger* Tu 4113. The single *m/z* expansion for the chromatogram is ± 5 ppm. ***b***, the Extracted Ion Chromatography (EIC) of *m/z* 1047.5863 $[M+Na]^+$ in wild-type *S. rapamycinicus* NRRL 5491 and *S. violaceusniger* Tu 4113. The single *m/z* expansion for the chromatogram is ± 5 ppm. ***c***, the LC-ESI-MS spectrum of **1** in wild-type *S. violaceusniger* Tu 4113. ***d***, the LC-ESI-MS spectrum of **1** in wild-type *S. rapamycinicus* NRRL 5491. ***e***, the LC-ESI-MS spectrum of **2** in wild-type *S. violaceusniger* Tu 4113. ***f***, the LC-ESI-MS spectrum of **2** in wild-type *S. rapamycinicus* NRRL 5491. ***g***, the LC-ESI-MS spectrum of **3** in wild-type *S. violaceusniger* Tu 4113. ***h***, the LC-ESI-MS spectrum of **3** in wild-type *S. rapamycinicus* NRRL 5491.

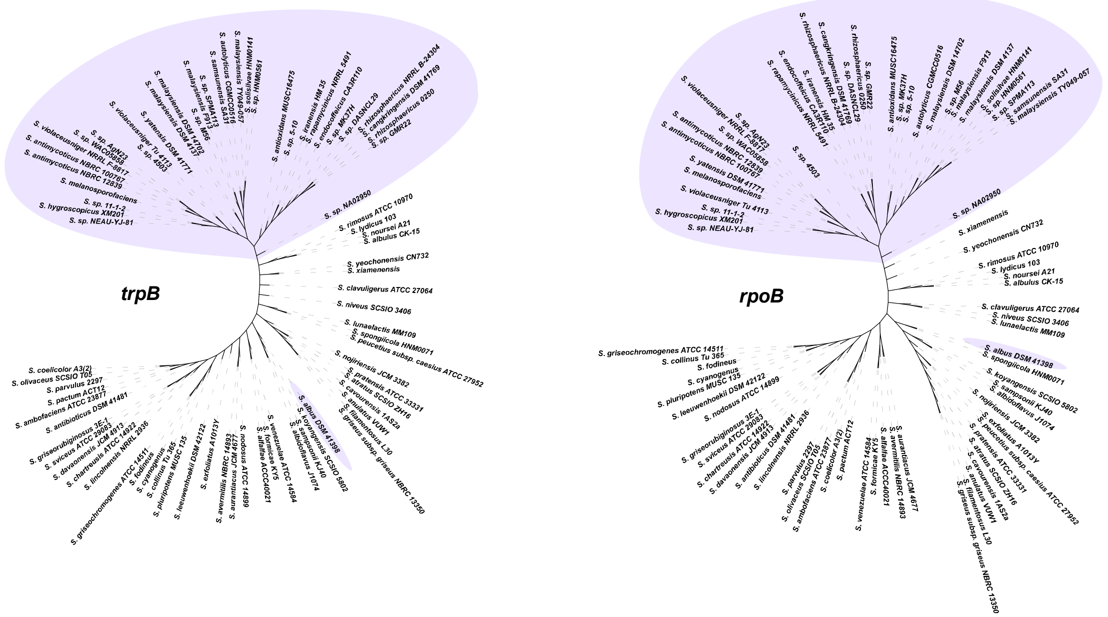


**Fig. 31**. The phylogenetic analysis of potential pteridic acids *Streptomyces* producers. The high-resolution *Streptomyces* sp. housekeeping genes: *trpB* (tryptophan synthase subunit beta) and *rpoB* (RNA polymerase subunit beta) were used in this analysis.

a

| | | 1 | 2 | 3 | 4 | 5 | 6 | 7 | 8 | 9 | 10 | 11 | 12 | 13 | 14 | 15 |
|---|---|---|---|---|---|---|---|---|---|---|---|---|---|---|---|---|
| *S. rapamycinus* NRRL 5491 | 1 | | 95.78 | 83.53 | 91.79 | 91.64 | 91.73 | 91.65 | 91.60 | 91.75 | 91.72 | 91.61 | 91.59 | 86.13 | 91.59 | 91.50 |
| *S. iranensis* HM 35 | 2 | 74.82 | | 83.59 | 91.80 | 91.72 | 91.77 | 91.68 | 91.62 | 91.79 | 91.63 | 91.62 | 91.64 | 86.10 | 91.73 | 91.59 |
| *S. albus* DSM 41398 | 3 | 10.75 | 10.61 | | 83.56 | 83.60 | 83.52 | 83.57 | 83.60 | 83.54 | 83.54 | 83.61 | 83.63 | 83.70 | 83.47 | 83.56 |
| *S. antimycoticus* NBRC 100767 | 4 | 70.38 | 68.49 | 11.36 | | 91.65 | 95.60 | 91.63 | 91.58 | 95.58 | 97.58 | 91.58 | 91.58 | 86.14 | 95.59 | 96.12 |
| *S. autolyticus* CGMCC0516 | 5 | 66.51 | 64.83 | 11.31 | 68.75 | | 91.64 | 98.77 | 98.91 | 91.71 | 91.52 | 98.92 | 98.80 | 86.07 | 91.52 | 91.43 |
| *S. hygroscopicus* XM201 | 6 | 66.63 | 65.41 | 11.15 | 73.49 | 69.52 | | 91.65 | 91.61 | 98.90 | 95.18 | 91.61 | 91.59 | 86.15 | 96.12 | 95.16 |
| *S. malaysiensis* DSM 4137 | 7 | 66.86 | 64.77 | 11.14 | 68.99 | 83.07 | 68.79 | | 98.80 | 91.70 | 91.48 | 98.81 | 98.87 | 86.05 | 91.52 | 91.41 |
| *S. solisilvae* HNM0141 | 8 | 65.56 | 62.94 | 10.79 | 67.61 | 87.21 | 67.84 | 89.60 | | 91.63 | 91.44 | 99.99 | 98.80 | 86.06 | 91.46 | 91.38 |
| *S.* sp. 11-1-2 | 9 | 67.91 | 66.17 | 11.21 | 74.30 | 70.01 | 87.13 | 70.25 | 69.05 | | 95.17 | 91.64 | 91.63 | 86.14 | 96.11 | 95.18 |
| *S.* sp. AgN23 | 10 | 68.18 | 64.40 | 11.33 | 78.86 | 67.16 | 70.61 | 67.94 | 66.24 | 72.65 | | 91.44 | 91.47 | 86.09 | 95.22 | 95.78 |
| S. sp. HNM0561 | 11 | 65.39 | 62.80 | 10.73 | 67.47 | 87.01 | 67.67 | 89.40 | 99.30 | 68.88 | 66.11 | | 98.80 | 86.07 | 91.46 | 91.39 |
| *S.* sp. M56 | 12 | 65.15 | 63.01 | 10.73 | 67.12 | 86.71 | 67.90 | 88.11 | 91.88 | 68.63 | 65.97 | 91.67 | | 86.08 | 91.47 | 91.37 |
| *S.* sp. NA02950 | 13 | 41.20 | 41.77 | 11.53 | 43.96 | 43.33 | 42.16 | 42.40 | 41.15 | 42.67 | 43.29 | 41.09 | 41.12 | | 86.11 | 86.19 |
| *S. violaceusniger* Tu 4113 | 14 | 67.65 | 67.66 | 11.52 | 76.23 | 68.53 | 73.87 | 68.27 | 66.21 | 75.44 | 72.44 | 66.08 | 66.28 | 43.88 | | 95.25 |
| *S. yatensis* DSM 41771 | 15 | 70.13 | 69.72 | 11.89 | 80.47 | 70.49 | 76.37 | 71.00 | 69.76 | 76.91 | 74.15 | 69.62 | 69.57 | 45.31 | 80.17 | |

**Upper:** average nucleotide identity **Lower:** alignment percentage

b

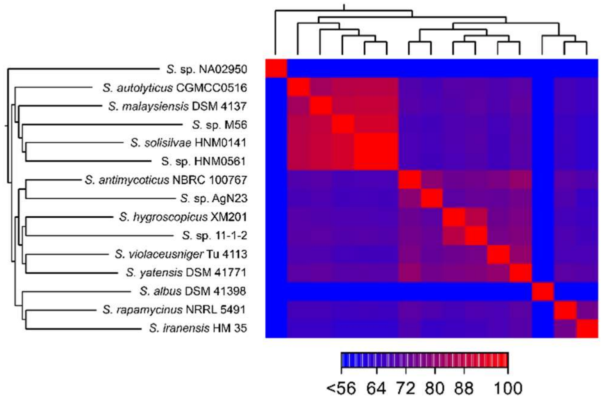


**Fig. 32**. Genome similarity analysis based on the alignment of 15 sequenced *pta*-containing *Streptomyces* genomes. ***a***, AP (alignment percentage) and ANI (average nucleotideidentity) values between genomes. The AP value represents the average percentage of aligned genomic regions between two genomes, whereas the ANI value stands for the percentage of exactly matching nucleotides for these aligned regions. ***b***, Heat map showing genetic similarity between genomes based on AP values.

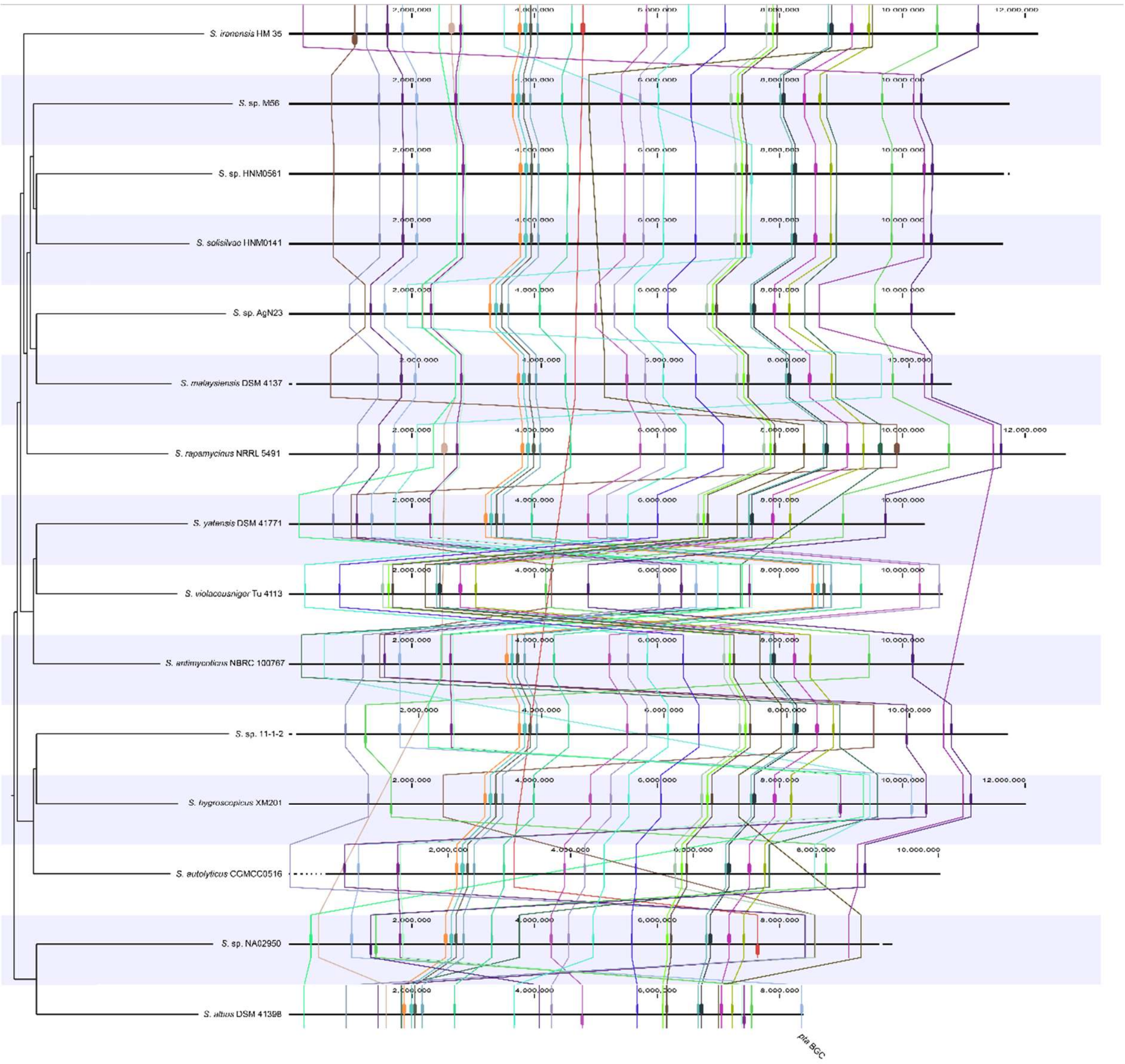


**Fig. 33**. The genome synteny analysis of selected 15 *pta*-containing *Streptomyces* strains. The colored squares each represent local alignment synteny blocks that are linked together among different genomes by lines with the same colors. The pteridic acids (pta) BGC is located in the end of *S. albus* DSM 41398 chromosome.

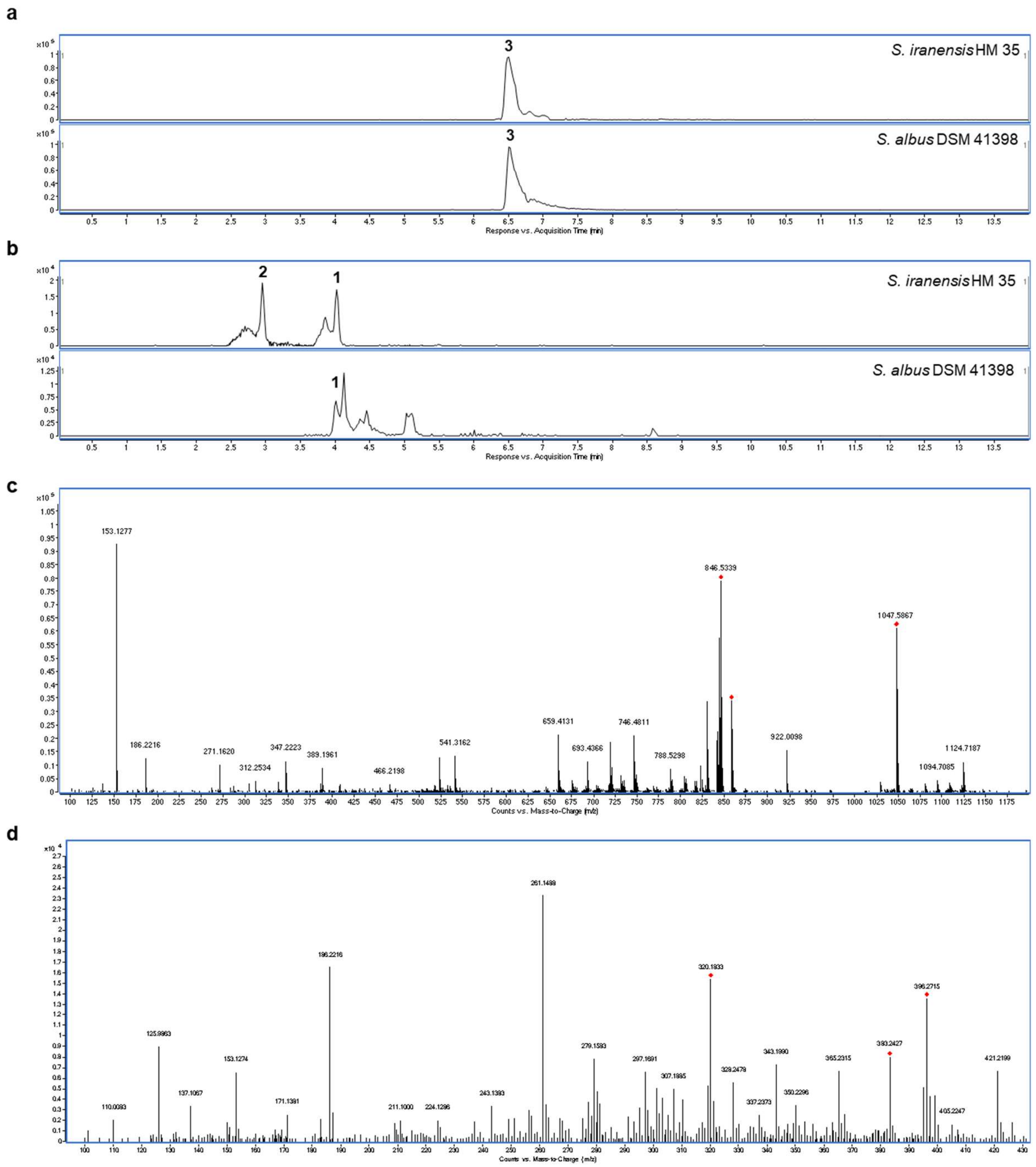


**Fig. 34**. The HPLC and mass spectrum analysis of metabolites in *S. albus* DSM 41398. ***a***, the Extracted Ion Chromatography (EIC) of *m/z* 383.2428 $[M+H]^+$ in wild-type *S. iranensis* HM 35 and wild-type *S. albus* DSM 41398. The single *m/z* expansion for the chromatogram is ± 5 ppm. ***b***, the Extracted Ion Chromatography (EIC) of *m/z* 1047.5863 $[M+Na]^+$ in wild-type *S. iranensis* HM 35 and wild-type *S. albus* DSM 41398. The single *m/z* expansion for the chromatogram is ± 5 ppm. ***c***, the LC-ESI-MS spectrum of **3** in wild-type *S. albus* DSM 41398. ***d***, the LC-ESI-MS spectrum of **1** in wild-type *S. albus* DSM 41398.

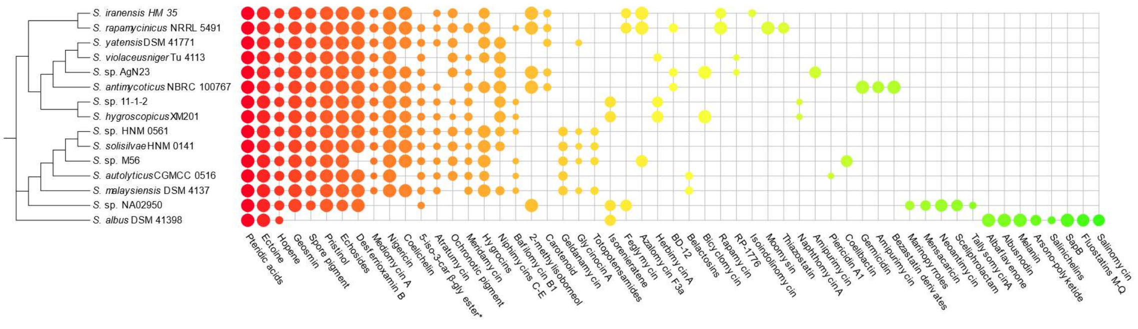


**Fig. 35**. The secondary metabolites biosynthetic gene clusters profiles of 15 *pta*-containing *Streptomyces* strains with complete genome information. BGCs with a similarity greater than 45% are displayed, and the size of the nodes represents the level of similarity *, 5-isoprenylindole-3-carboxylate β-glycosyl ester.

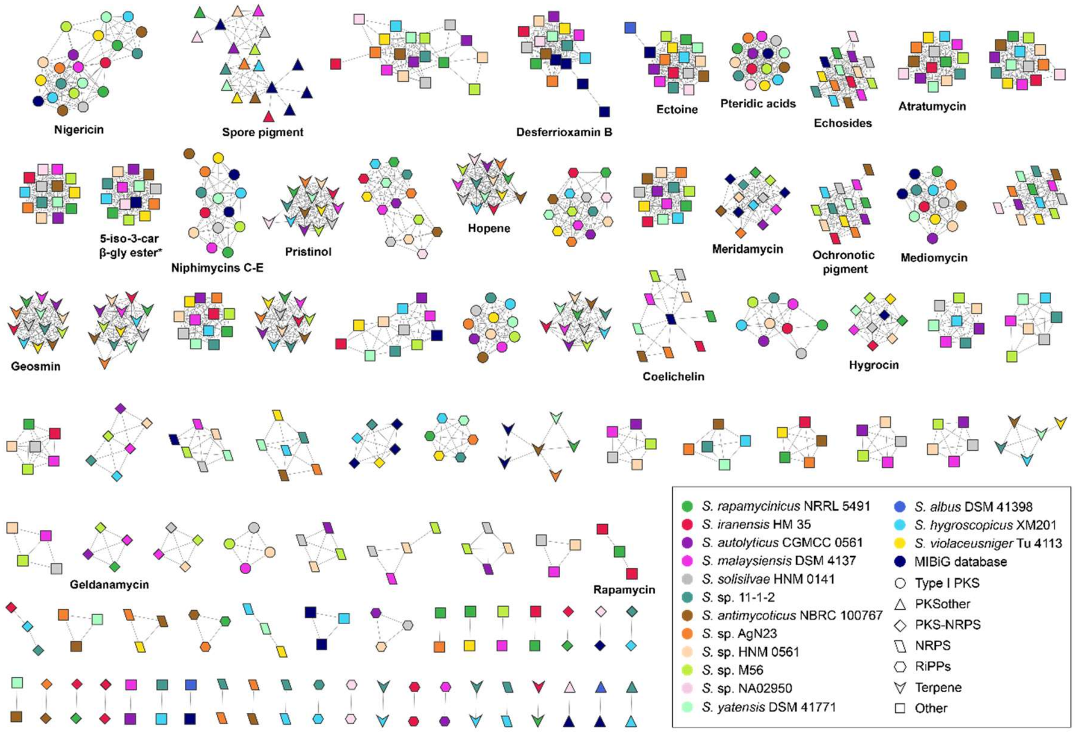


**Fig. 36**. The secondary metabolites BGCs similarity network of 15 pta-containing *Streptomyces* strains with complete genome information. Different strains and types of secondary metabolites are represented by different colours and shapes, respectively. *, 5-isoprenylindole-3-carboxylate β-glycosyl ester.
